# Independent Genomic Sources of Brain Structure and Function

**DOI:** 10.1101/2021.01.06.425535

**Authors:** Sourena Soheili-Nezhad, Christian F. Beckmann, Emma Sprooten

## Abstract

**Introduction:** The last decade has seen a surge in well powered genome-wide association studies (GWASs) of complex behavioural traits, disorders, and more recently, of brain structural and functional neuroimaging features. However, the extreme polygenicity of these complex traits makes it difficult to translate the GWAS signal into mechanistic biological insights. We postulate that the covariance of SNP-effects across many brain features, as be captured by latent genomic *components* of SNP effect sizes. These may partly reflect the concerted multi-locus genomic effects through known molecular pathways and protein-protein interactions. Here, we test the feasibility of a new data-driven method to derive such latent components of genome-wide effects on more than thousand neuroimaging derived traits, and investigate their utility in interpreting the complex biological processes that shape the GWAS signal.

**Methods:** We downloaded the GWAS summary statistics of 3,143 brain imaging-derived phenotypes (IDPs) from the UK Biobank, provided by the Oxford Brain Imaging Genetics (BIG) Server (Elliott et al. 2018). Probabilistic independent component analysis (ICA) was used to extract two hundred independent genomic components from the matrix of SNP-effect sizes. We qualitatively describe the distribution of the latent component’s loadings in the neuroimaging and the genomic dimensions. Gene-wide statistics were calculated for each genomic component. We tested the genomic component’s enrichment for molecular pathways using MSigDB, and for single-cell RNA-sequencing of adult and foetal brain cells.

**Results:** 200 components explained 80% of the variance in SNP-effects sizes. Each MRI modality and data processing method projected the imaging data into a clearly distinct cluster in the genomic component embedded space. Among the 200 genomic components, 157 were clearly driven by a single locus, while 39 were highly polygenic. Together, these 39 components were significantly enriched for 2,274 MSigDB gene sets (fully corrected for multiple testing across gene-sets and components). Several components were sensitive to molecular pathways, single cell expression profiles, and brain traits in patterns consistent with knowledge across these biological levels. To illustrate this, we highlight a component that implicated axonal regeneration pathways, which was specifically enriched for gene expression in oligodendrocyte precursors, microglia and astrocytes, and loaded highly on white matter neuroimaging traits. We highlight a second component that implicated synaptic function and neuron projection organization pathways that was specifically enriched for neuronal cell transcriptomes.

**Conclusion:** We propose genomic ICA as a new method to identify latent genetic factors influencing brain structure and function by multimodal MRI. The derived latent genomic dimensions are highly sensitive to known molecular pathways and cell-specific gene expression profiles. Genomic ICA may help to disentangle the many different biological routes by which the genome defines the inter-individual variation of the brain. Future research is aimed at using this method to profile individual subjects’ genomic data along the new latent dimensions and evaluating the utility of these dimensions in stratifying heterogeneous patient populations.

## 1. Introduction

Increasingly large genome-wide association studies (GWAS) show that brain disorders and related traits are driven by polygenic factors. Polygenicity has long been assumed for complex behavioural traits, but it is only now that we have come to fully appreciate the extreme form this polygenicity takes^1^. The brain is the logical organ of convergence for genetic effects on behavioural phenotypes, and quantitative neuroimaging can putatively trace some of these effects in living humans non-invasively^2^. Therefore, understanding the genetic underpinnings of neuroimaging traits, or endophenotypes^3^, is thought to give important insights into the causal processes driving variations of human behaviour in a healthy or disease population. However, two important caveats have prevented fulfilling this promise so far. Firstly, the dimensionality of the imaging and genetic data easily approaches millions of genetic variants and thousands of brain imaging features. Thus, the combination of neuroimaging and genetics aggravates the problems of multiple comparisons and interpretability that already exist within both fields of research. Secondly, the small effect size of each genetic variant, usually single-nucleotide polymorphisms (SNPs), complicates the translational potential of a GWAS and pharmacological targeting, even though such genomic effects may enlighten some aspects of the underlying pathways driving a complex polygenic trait^4–6^. In this paper, we present a new method to reduce the enormous genome-wide and brain-wide data into a lower dimensional space and demonstrate that it enables drawing a more interpretable picture of the many ways by which millions of genomic variants collectively influence the brain. Moreover, this method projects the genome-wide data into new latent factors that are readily applicable to individual data, in a way that can help describe biological heterogeneity in the context of both population-wide summary statistics and individual subjects’ raw genotypes. Ultimately, this approach is geared to facilitate moving from descriptive neuroimaging-genome associations towards personalised medicine.

Theoretically, the brain lies intermediate between the causal chain of events from genes to behaviour. While the brain’s genetic architecture, due to its proximity to cellular and molecular mechanisms, does seem to be less complex and less polygenic than the behavioural phenotypes, it still is marked by the aggregated effects of many variants with small effects^7,8^. Therefore, translation of recent GWAS of brain imaging traits and to causal mechanisms is nearly as complicated as those of the clinical phenotypes in psychiatry and related behavioural traits. To tackle this problem, propagating SNP effects on the brain into latent factors that are biologically coherent may be helpful. Such latent factors, may give better insight into possible mechanisms of action, and into the heterogeneous basis of these mechanisms in the future. In the current paper, we refer to these multi-SNP latent factors as genomic *components.* Here, we present a new method to derive these components, aimed to address the above challenges of interpretability and clinical translation of univariate GWAS.

Several types of polymorphisms underpin variability of the human genome across the population, such as SNPs, short insertion/deletions, and long structural variations. In GWAS, usually the association of SNPs - the most abundant type of DNA variation - is assessed with a trait of interest using a univariate regression model. In a typical scenario, a SNP-phenotype association may expose the effect of a gene (or nearby genes) on the trait. However, the mechanistic influence of a putative causal gene on a polygenic trait is usually not univariate, but rather takes place in concert with other (potentially distant) causal variants, particularly when they collectively tag multiple genes of a coherent molecular pathway. It stands to reason that such functional synchronicity between distant genes could partly be reflected in the covariance of SNP effect sizes across multiple biological traits, such as those extracted from high-dimensional imaging data. That is, for brain imaging traits, some groups of SNPs, regardless of their genomic distance, may collectively share their patterns of effect sizes across different brain features extracted from different brain regions and/or imaging modalities.

Shared effects of SNPs on different neuroimaging traits have indeed been observed by genetic correlation analyses, including features extracted from subcortical structures or the cerebral cortex^9,10^. In this regard, the interregional pattern of effects of two SNPs that lie in two biologically distant pathways may be more orthogonal and independent than those that lie in similar pathways. Here, we show that it is possible and informative to extract statistically independent latent features from genome-wide SNP effect size distributions in a data-driven way, particularly now that GWAS summary statistics on thousands of traits are available, owing to contemporary ‘biobank scale’ data^11^.

Magnetic resonance imaging (MRI) allows us to measure different aspects of local and global brain structure and function via different image acquisition pulse sequences (a.k.a. MRI “modalities”) and analysis techniques. Each MRI modality informs on some aspect of brain morphology and function in a spatially defined coordinate of regional anatomy. A GWAS of a neuroimaging feature provides information on which SNPs are associated with that particular feature, and via further bioinformatics analyses, may also give insight into underlying biological mechanisms underpinning this association. However, polygenicity limits the sensitivity and specificity of the univariate GWAS signal to molecular pathways and gene sets. Here, we aimed to investigate if a hidden structure exists among the summary statistics across brain regions and MRI modalities. We seek to find out if an unsupervised source separation method, independent component analysis (ICA), captures some multigenic factors that are biologically more coherent and interpretable than the mass univariate SNP associations. ICA is a computational source separation method for decomposing a linearly mixed signal into its constituents. ICA identifies statistically independent sources that drive a mixed signal as observed from different angles, here the angles being different brain regions and neuroimaging features, and the hidden sources being multigenic drives of their variability. In the current application, we use ICA for the first time to project a big array of GWAS data into a lower dimensional space, arriving at genomic ‘sources’ (hitherto referred to as genomic ‘components’) that we show may improve the sensitivity of the GWAS to molecular pathways, and thereby facilitate their interpretation in terms of plausible biological mechanisms. In addition, we explore the degree of specificity of the derived genomic components to various imaging modalities of the brain MRI traits.

## 2. Methods

### 2.1. Genomic ICA

We downloaded the GWAS summary statistics of 3,143 brain traits from the UK Biobank, provided by the Oxford Brain Imaging Genetics (BIG) Server (http://big.stats.ox.ac.uk; based on^10^). In brief, Elliott et al. used the processed data of the first wave of the UK biobank cohort (n=8,428 subjects). Multimodal MRI acquisitions included structural, functional, diffusion, fluid-attenuated inversion recovery (FLAIR) and susceptibility-weighted scans. Each of the scans was processed as detailed in^10^, eventually yielding 3,143 brain imaging-derived phenotypes (IDPs), which were the continuous brain straits entered into GWAS^10^. We concatenated all the SNP effect sizes (i.e. GWAS regression betas) across all the IDPs to create a brain-wide and genome-wide IDPxSNP matrix. In this matrix, we excluded those IDPs (i.e. rows) encoding the strength of node-to-node functional connectivity, as they have been previously shown to possess low heritability^10^. Among SNPs, only variants with high cross-trait variance (above the 95th percentile) were kept in the analysis, as we observed that by focusing the model to this subset of high-variance SNPs the component weights are regularized to follow a Gaussian distribution with minimal kurtosis. This is a useful property and facilitates the subsequent separation of the multivariate ‘genetic’ signal from the background Gaussian noise by parametric fitting and z-transformation. In this regard, variance-based filtering is still expected to keep potentially interesting SNPs in the model, as ICA inherently relies on cross-trait variance for decomposing the latent components by optimizing a metric of statistical independence such as neg-entropy^12^. This procedure left a matrix of 1,448 IDPs by 534,063 high-variance SNPs for further analysis (Figure 1).

**Figure 1.**
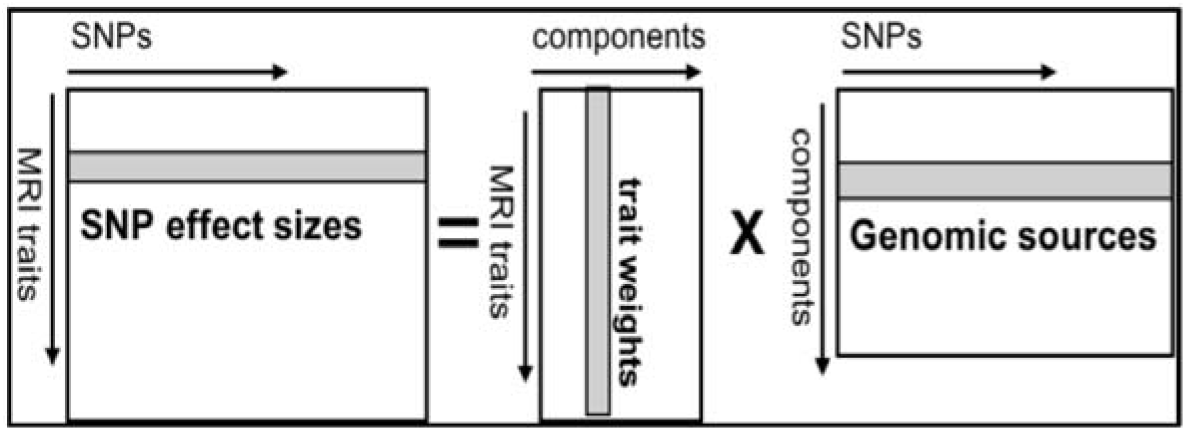
SNP effect by MRI trait ICA decomposition. On the left side of the equation are is the input data, consisting of GWAS output. On the right are the 200 resulting genomic sources (or components) with SNP-loadings for each source, together with the respecitive arrays of MRI components, which consist of weights for each trait on each component.

Probabilistic ICA, MELODIC^13^, was used to identify independent genomic sources of this IDPxSNP matrix. The mean and the variance of the SNP effects were not removed/normalized prior to the decomposition, as they bear biological information in GWAS, contrary to ICA in other modalities such as functional neuroimaging^13^. Two hundred *independent genomic components* were extracted, each of which consists of a latent genomic factor (in the SNP-dimension) of SNP weights and a corresponding IDP-loading vector (in the MRI dimension) indicating the degree to which each IDP is associated with that genomic factor/component (Figure 1). ICA allows for different genomic components to overlap in the continuous space of the genome, i.e SNPs can have similarly high or low weights in multiple sources, and thus provides a ‘soft’ clustering of the SNPs based on their statistical independence in driving brain traits.

### 2.2. Gene-set enrichment of the genomic sources for Gene Ontology terms and single cell gene-expression

A Gaussian distribution was fit to the SNP-weights of each genomic component, to identify SNPs with the strongest contribution per component. SNP-wise z-scores were then derived from this Gaussian distribution and transformed to P-values. Using hg19 genomic coordinates and gene-wide tests, the top-SNP model of MAGMA^14^ (version 1.08; https://ctg.cncr.nl/software/magma) was applied to calculate gene-wide statistics for each component. Thereafter, each genomic component was tested for the enrichment of gene ontology (GO) terms using MAGMA’s gene-set enrichment test. These gene-sets were retrieved from the Molecular Signatures Database^15^ (MSigDB; https://www.gsea-msigdb.org). To avoid gene-set enrichments being driven by a single, highly significant locus, we excluded gene sets with fewer than 5 genes and clear, single-locus components (see Figure 2 and section 3.1. below) from this analysis.

**Figure 2.**
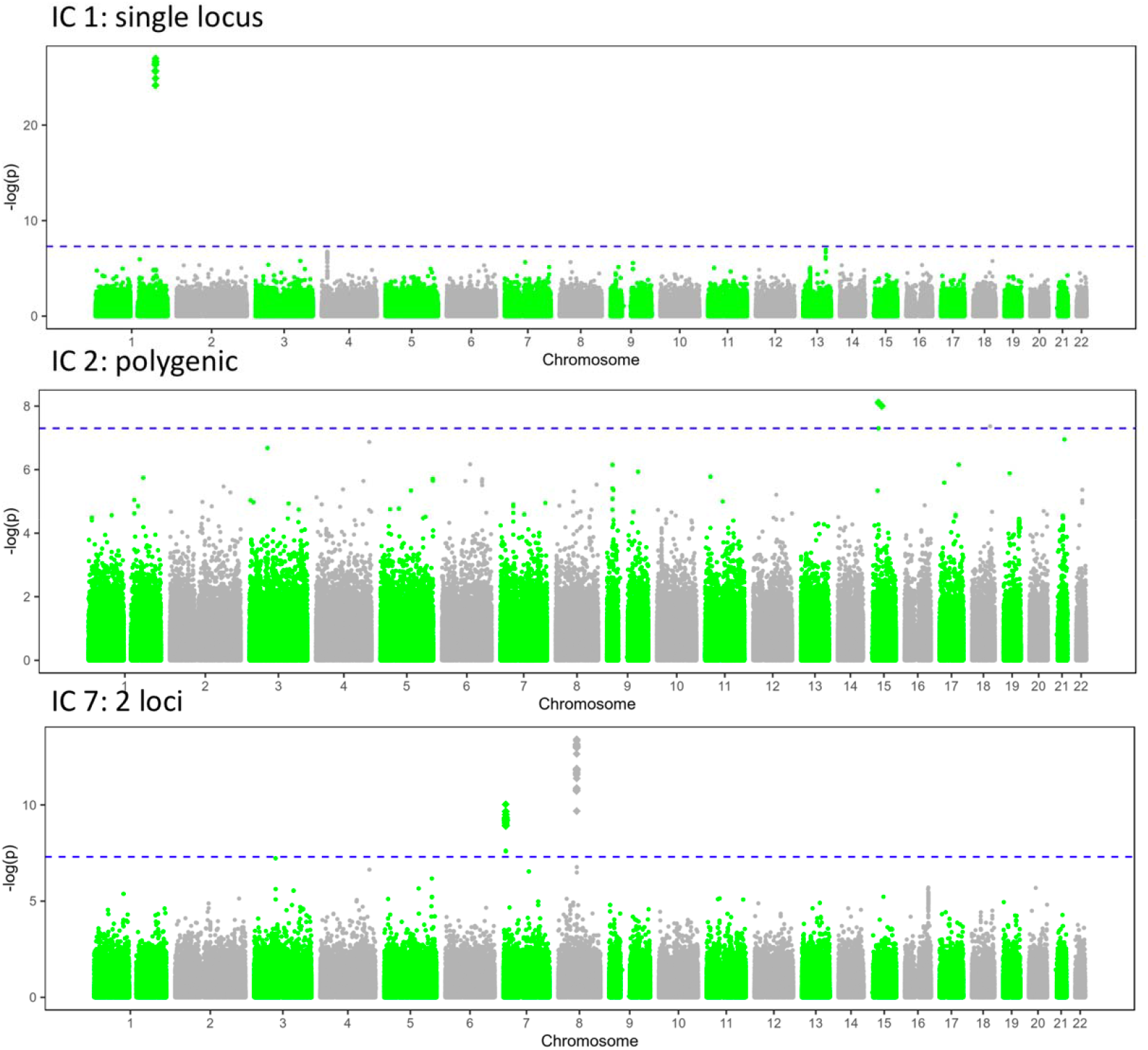
First of each “type” of component: IC1 was classified as single locus component, IC2 as polygenic and IC 7 as 2 locus-component.

In addition, cell type specificity of 3 highlighted genomic components was assessed using MAGMA’s gene property analysis^16^ and the single cell gene expression profiles of developmental and adult human brain (iPsych RNA-seq; downloaded from https://github.com/Kyoko-wtnb/FUMA-webapp^17^). Twenty-six cell types in the adult brain and 36 cell types in the developmental brain tissue database were tested for enrichment. To control the false positive rate, we applied FDR-correction across all cell types.

### 2.3. Modality-specific genomic components

Each MRI modality (structural, functional or diffusion) informs on a particular aspect of neurobiology, and different MRI features may have common or distinctive cellular and genomic underpinnings^18^. To identify genomic components specific to MRI modalities, we used L1-regularized logistic regression (LASSO). The modality classes of interest included cortical thickness (n= 212 IDPs), cortical surface area (n=214 IDPs), diffusion MRI measures (n= 675 IDPs), amplitude of the regional resting-state signal fluctuation (n=76 IDPs), and the volume of cortical and subcortical brain regions (n= 227 IDPs). To further visualize similarities and clustering of the IDPs with regard to the genomic drives, IDP loadings were transferred to a two dimensional embedded space using t-distributed stochastic neighbour embedding (t-SNE^19^).

## 3. Results

### 3.1. Description of the genomic components

The 200 independent genomic components explained 80% of the variance in SNP-effects across the IDPs. Upon visual inspection of the Manhattan plots of the genomic sources, 157 out of the 200 components were clearly driven by a single locus. In contrast, 39 components were polygenic in nature, with no clearly identifiable peak. Another 4 components had 2 discernible peaks. All Manhattan plots are available in Supplementary figure 1.

Figure 2 shows the first three components of each type for illustration. The full dataset of SNP-weights for all 200 components will be made publicly available after the final version of this manuscript is published.

The 157 components that were clearly driven by a single locus collectively implicated 67 intronic and 2 exonic top variants mapping to gene transcription regions. The four double locus components further implicated five intronic variants. These components and their top significant loci and closest genes are listed in Supplementary table 1.

### 3.2. Modality specificity of genomic components

t-SNE showed a clear clustering pattern for IDPs extracted from similar modalities, including structural, diffusion, functional and susceptibility-weighted MRI proximity of MRI-derived phenotypes in the embedded space indicates similar genetic influences for traits within each imaging modality, and different genetic influences on traits across modalities (Figure 3). We did not detect cross-modal regional specificity. Different methods used for extracting morphological features from T1-weighted images including regional brain volume, cortical surface area and thickness resulted in IDPs mapping to separate genomic clusters. In contrast, the two methods used for diffusion MRI processing, TBSS and tractography, showed similar genetic influences.

**Figure 3.**
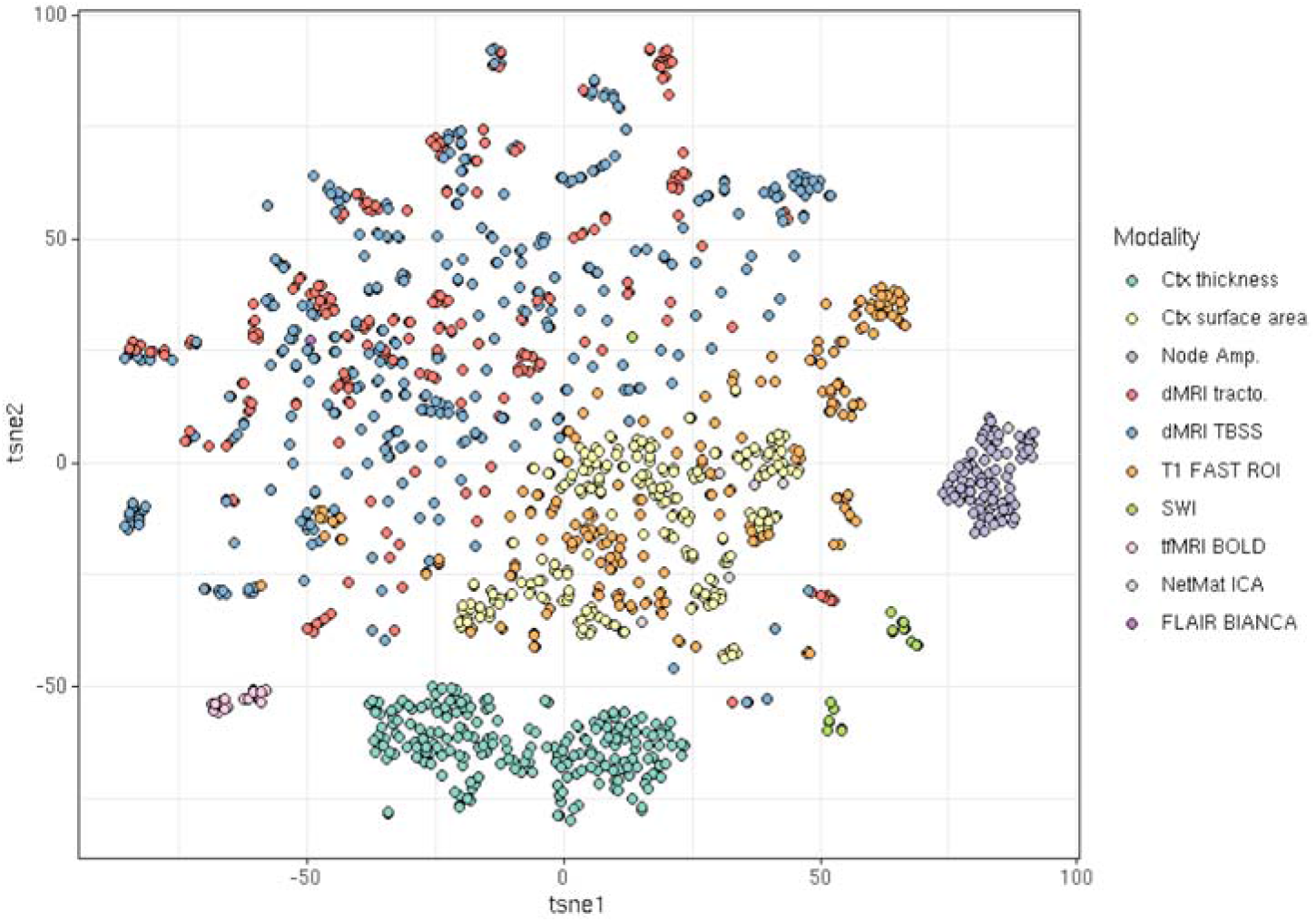
t-SNE embedding of IDP loadings in the genomic ICA space

Following logistic regression, we found that different subsets of the 200 genomic components predicted IDPs extracted from each modality. Notably, component #86 (highlighted in detail below) demonstrated the highest contribution in classifying both diffusion MRI and cortical surface area (Supplementary table 3). This component was the only one with strong predictive power in more than one imaging modality. All classification accuracies were above 98% in a 100-fold cross-validation test (Supplementary figure 2). Supplementary table 3 contains all the statistics/regression betas for each component across the modalities.

### 3.3. ene-set enrichment of multi-locus components

The 39 polygenic components extracted by ICA were enriched for 2,274 MSigDB gene sets related to different biological processes (FDR-corrected across all gene-sets and 39 components, top 20 terms in table 1 and the full enrichment in Supplementary table 2).

**Table 1:** Top Enriched pathways across all 39 polygenic components

| IC# | GO term | p (FDR) |
| --- | --- | --- |
| 199 | autophagy of nucleus | 5.75E-28 |
| 200 | Paraspeckles | 4.94E-20 |
| 200 | positive regulation of hematopoietic progenitor cell differentiation | 2.97E-18 |
| 133 | de novo imp biosynthetic process | 1.35E-13 |
| 86 | extrinsic component of presynaptic membrane | 1.53E-13 |
| 199 | ATG1 ULK1 kinase complex | 4.60E-13 |
| 200 | regulation of lymphoid progenitor cell differentiation | 4.90E-13 |
| 86 | peptide proton symporter activity | 2.49E-12 |
| 111 | lamin filament | 3.77E-12 |
| 103 | methylglyoxal metabolic process | 1.08E-11 |
| 99 | intraciliary transport particle B binding | 5.74E-11 |
| 100 | negative regulation of apoptotic process in bone marrow cell | 1.56E-10 |
| 6 | citrate metabolic process | 1.61E-10 |
| 7 | actin filament based transport | 6.69E-10 |
| 100 | regulation of myeloid dendritic cell activation | 7.42E-10 |
| 86 | annulate lamellae | 7.93E-10 |
| 145 | transmembrane receptor protein tyrosine phosphatase signaling pathway | 1.10E-09 |
| 86 | postsynaptic actin cytoskeleton | 1.12E-09 |
| 95 | purine nucleobase transport | 1.30E-09 |
| 86 | regulation of synapse structural plasticity | 1.43E-09 |

### 3.4. Highlighted component #54: axonal regeneration

One of the polygenic components (IC #54) showed highly significant gene-set enrichment for multiple pathways related to axon regeneration (table 2). In adult brain cells, component 54 was specifically and significantly enriched for gene expression in oligodendrocyte precursors, microglia and astrocytes (table 3). This component also showed nominal enrichment for average gene expression in developing brain cells (P=0.048), with the most enrichment for expression in oligodendrocyte precursors, but this was not significant after FDR-correction (P=0.003; P_FDR=0.058; table 3). Fitting with a role of axon regeneration, diffusion weighted imaging traits weighted highly on this component, as did cortical surface area traits (Supplementary table 3).

**Table 2:**
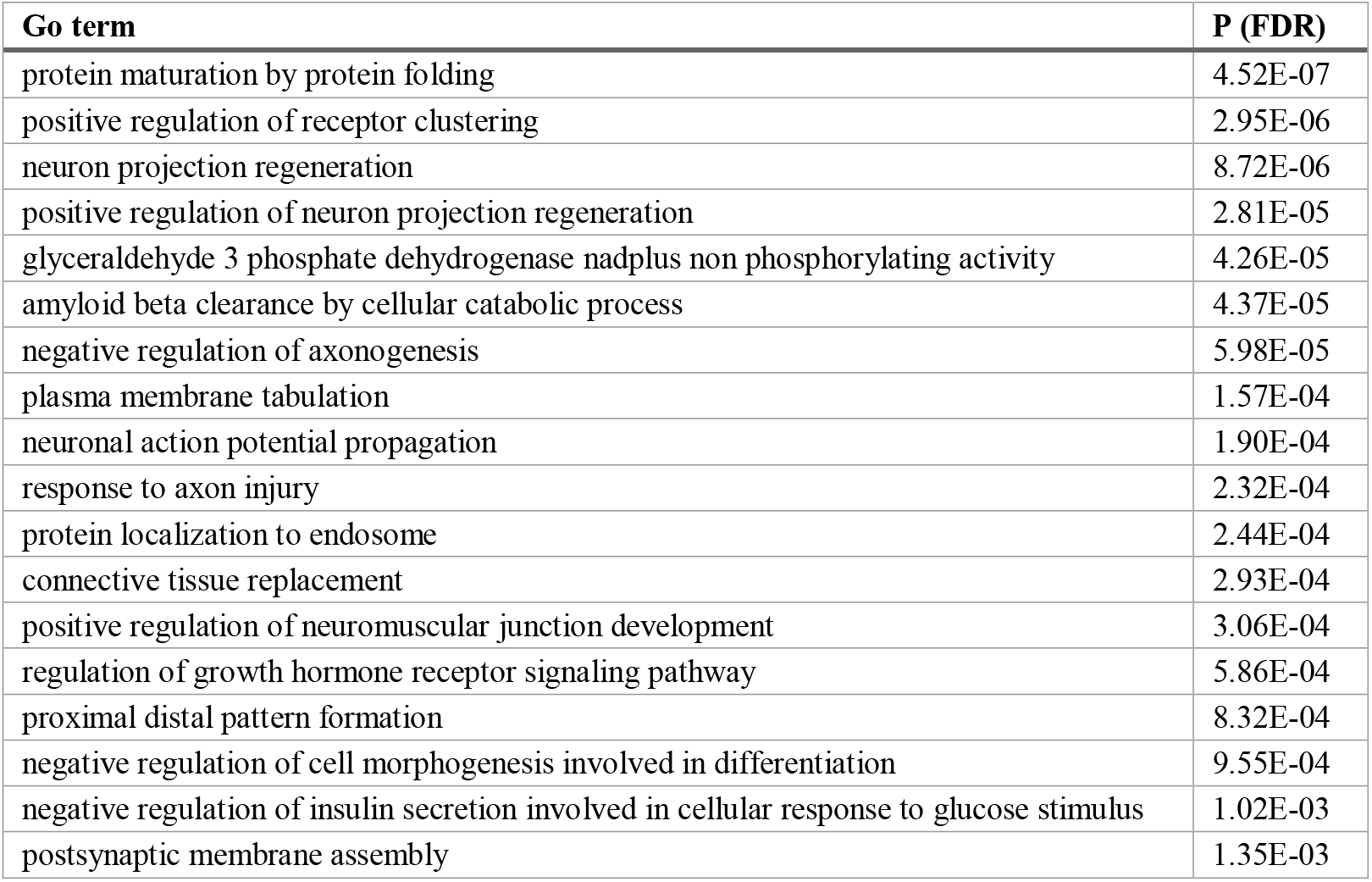
Signifcantly enriched MSigDB gene sets for Highlighted Component 54, with multiple significantly enriched gene-sets indicating a role for Axon Regeneration. P-values are FDR-corrected across gene-sets and the 39 polygenic components.

**Table 3:** Enrichment of Highlighted Component 54 for different cell types in developing and adult brain tissue. Results were only significant for OPC’s, microglia and astrocytes in the adult brain, but for better illustration all results with uncorrected P < 0.05 are shown.

| Developing brain | P | P (FDR) | Adult brain | P | P (FDR) |
| --- | --- | --- | --- | --- | --- |
| Average | 0.048 | - | Average | 0.163 | - |
| Fetal OPC | 0.003 | 0.058 | OPC | 0.002 | 0.023 |
| Ex6 | 0.006 | 0.058 | Microglia | 0.003 | 0.023 |
| Ex8 | 0.006 | 0.058 | Astro | 0.003 | 0.023 |
| In5 | 0.006 | 0.058 | Ex6b | 0.022 | 0.145 |
| Ex7 | 0.01 | 0.06 | Per | 0.028 | 0.145 |
| Ex1 | 0.01 | 0.06 | Oligo | 0.05 | 0.216 |
| In8 | 0.012 | 0.064 |  |  |  |
| Ex2 | 0.019 | 0.08 |  |  |  |
| In6 | 0.02 | 0.08 |  |  |  |
| Fetal astrocytes | 0.023 | 0.083 |  |  |  |
| In7 | 0.03 | 0.097 |  |  |  |
| In3 | 0.035 | 0.099 |  |  |  |
| Ex3 | 0.037 | 0.099 |  |  |  |
| Ex5 | 0.039 | 0.099 |  |  |  |
| In1 | 0.041 | 0.099 |  |  |  |

### 3.5. Highlighted component #86: synaptic function

Component 86 is of special interest because it best distinguished different modalities, in particular having opposite weights on surface area and DWI-derived traits (Section 3.2, Supplementary table 3). GO term enrichment of this component implicated synaptic function, cytoskeletal plasticity and the extracellular matrix (table 4). Gene property test using single cell transcriptome profiles showed that this component was mostly representative of the excitatory neurons (table 5).

**Table 4.**
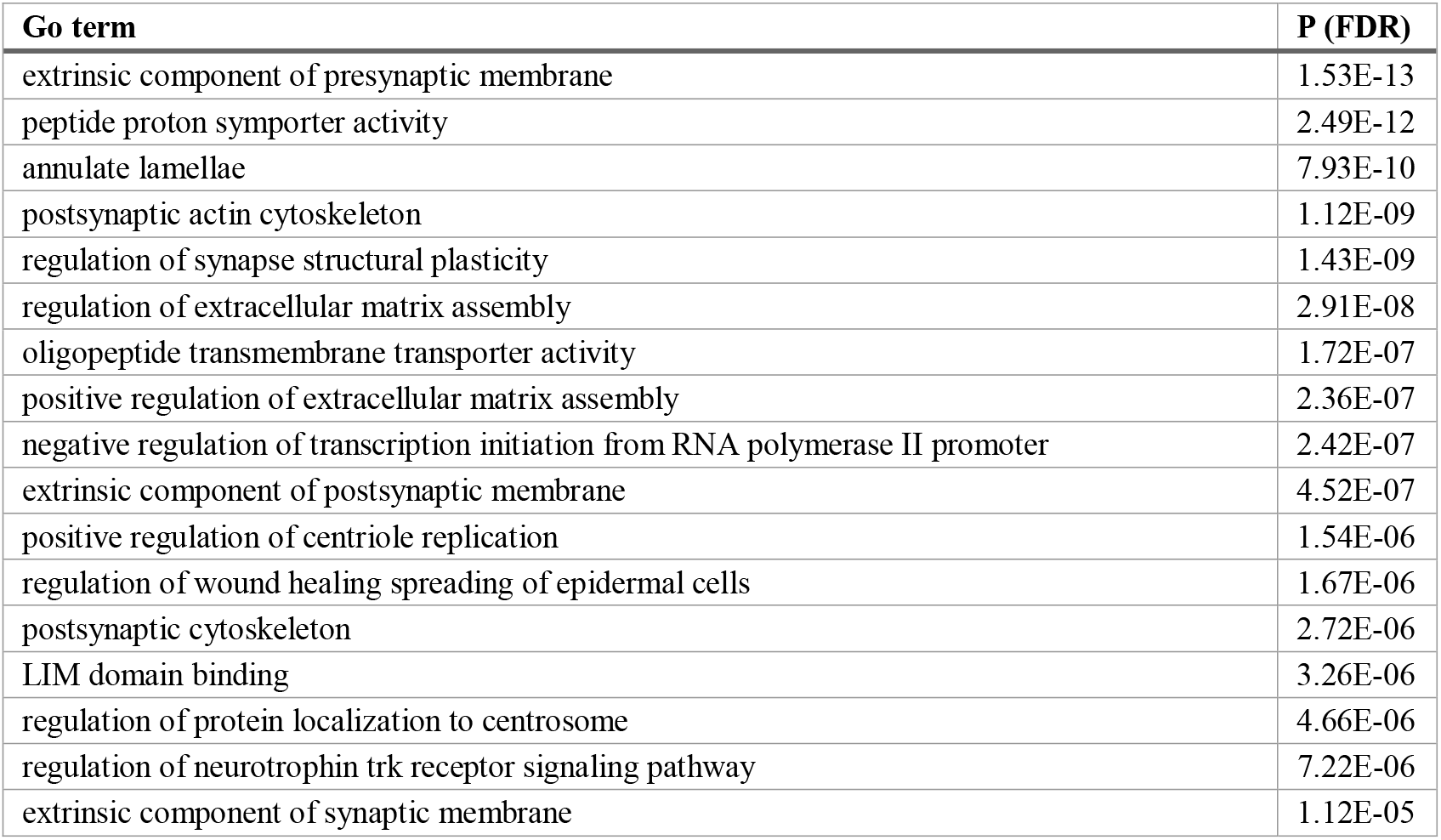
Signifcantly enriched MSigDB gene sets for Highlighted Component 86, with multiple significantly enriched gene-sets indicating a role for synaptic function. P-values are FDR-corrected across gene-sets and Bonferroni-corrected across the 39 polygenic components.

**Table 5.**
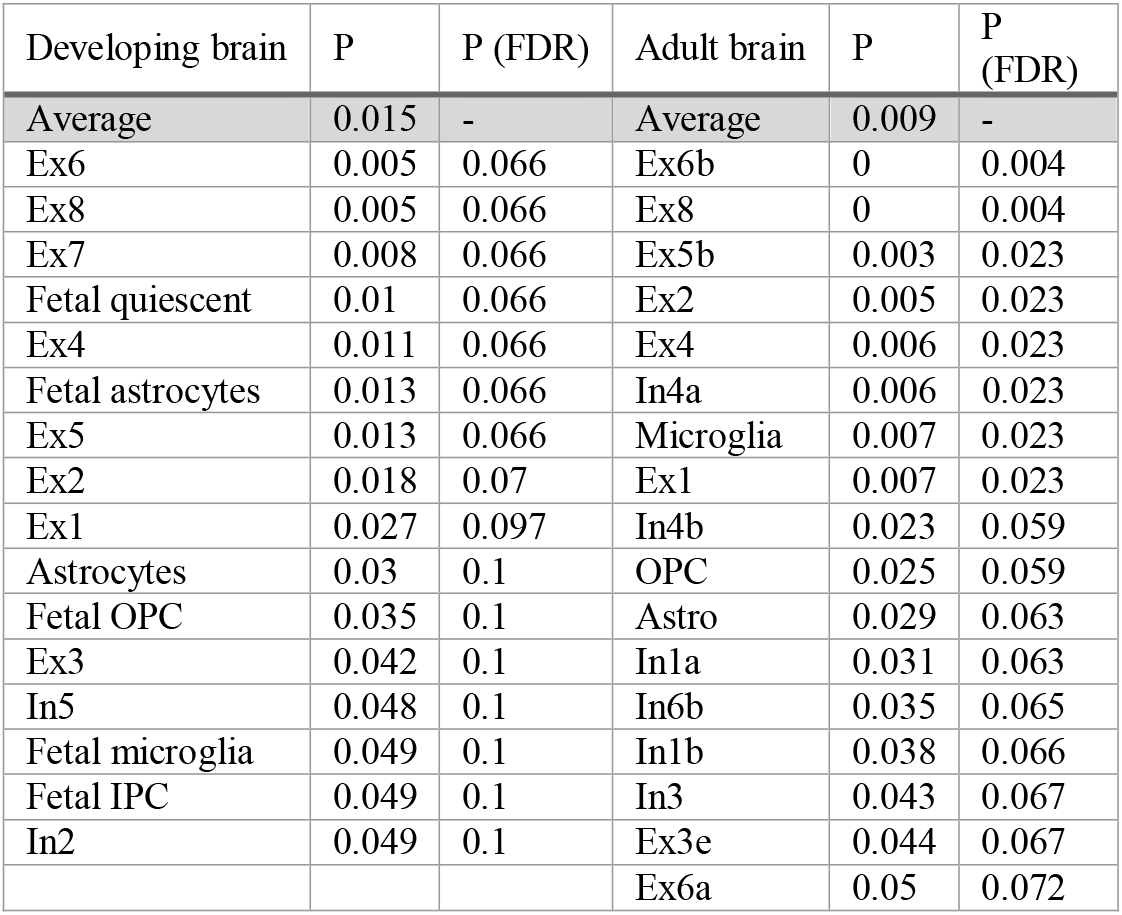
Enrichment of Highlighted Component 86 for different cell types in developing and adult brain tissue. Results were most significant for excitatory neurons in the adult brain, but for illustration all results with uncorrected P < 0.05 are shown.

### Highlighted component #181: single-locus protocadherin-A

Component #181, one of the components with a single strong locus, mapped to the 5q31 region where a dense cluster of protocadherin genes are located, These genes code for several members of the neural cadherin-like family genes (Figure 4), with roles in synapse-to-synapse adhesion and contact. A total of 137 variants in high LD (lead SNP: rs181578390, intronic, z-score=13.95) contributed to the multivariate signal of this component, among which 17 were exonic (top SNP: rs201493000, missense: PCDHA11:Q61>P). Interestingly, the large number of protocadherin genes (α=15, β=16 and γ=22) in this cluster is stochastically expressed in neurons due to alternative promoter choices. This randomness results in an enormous variability of tetramer combinations for these neural adhesion molecules, a feature that may generate single cell diversity of neurons and provide an address code for synaptic connectivity^20^. The protocadherin synaptic proteins are associated with various neuropsychiatric disorders^21^ and their genomic component mostly affected diffusion and cortical surface area phenotypes (Supplementary table 3).

**Figure 4.**
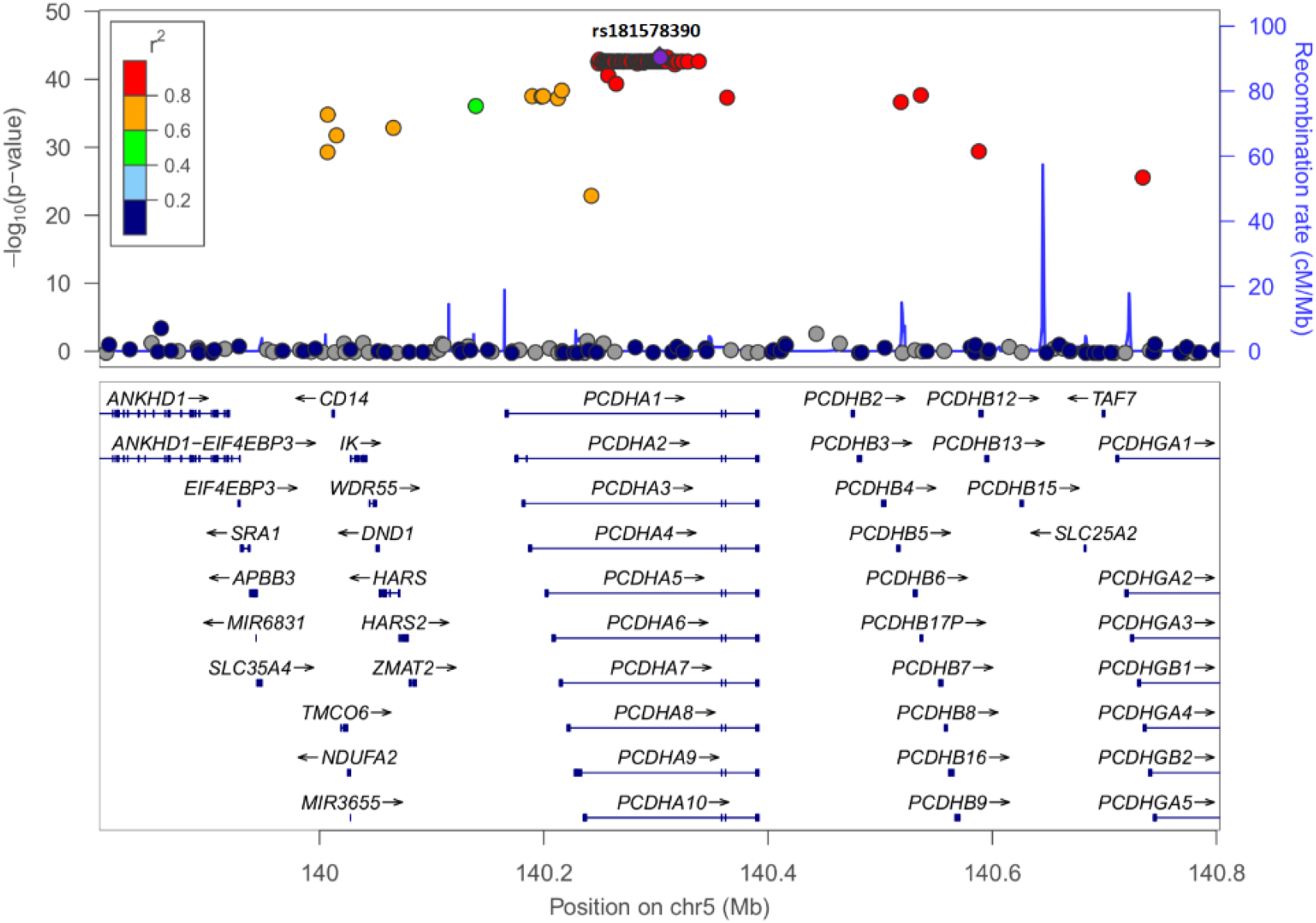
Protocadherin locus (#181)

## 4. Discussion

The complex, extremely polygenic architecture of psychiatric, behavioural, and brain traits has so far complicated the translational impact of GWAS in terms of concrete mechanistic insights and direct clinical impact. The rapidly expanding collection of GWAS statistics can now be seen as a source of ‘raw’ data in itself that can be better interpreted after further modelling and post-hoc analysis. We present for the first time a fully data-driven, multivariate method towards this goal using GWAS summary statistics. We demonstrate the sensitivity and specificity of this method for potentially interesting gene ontologies and functions. The preliminary results seem to inform on the molecular origins of brain variability and their multivariate genetic architecture. Particularly, the results show that genomic ICA may improve the interpretability and translational impact of GWAS, although this needs to be further validated and expanded in independent cohorts. GWAS is better interpreted when complemented with other high-throughput data such as transcriptome-wide association studies (TWAS), epigenomics and proteomics, all of which provide different renditions of the same underlying biology (i.e. latent factors) but from different angles. For this aim, ICA has been historically used in other disciplines as a linear source separation or multimodal data fusion method^22^. This makes ICA a suitable candidate for combining genomic and transcriptomic data across modalities in the future as well.

Genome-wide pathway analysis is a useful method that has been widely applied to identify biological mechanisms underlying polygenic disorders^6^. Molecular pathway enrichment analyses of raw GWAS statistics tend to identify many pathways, each of with only modest enrichment, despite sample sizes of N>100,000. Univariate pathway enrichments do not model polygenic interactions and concerted, pleiotropic effects in the genome-wide data. For psychiatric disorders, these pathways range from neurogenesis and synaptic signalling processes in schizophrenia^23^, motor coordination, regulation of insulin secretion, and endocannabinoid signaling in bipolar disorder^24^ and synaptic structure and behavioural modulation in major depressive disorder^25^. Moreover, the causal pathways driving gene-brain associations include complex gene-environment interplay and are partly mediated via genetic effects on environmental factors as well^26^. Given the number of causal routes from gene to brain to psychopathology this permits, it should be no surprise that univariate GWAS findings are extremely polygenic, and specific neither to molecular pathways, nor to brain regions or tissues. We suggest that ICA may partly mitigate this problem by segregating the large array of SNP effects into a reduced number of independent genomic sources across thousands of GWASs, each one with a statistically independent distribution. This is particularly useful for a complex and hierarchically organized organ such as the human brain and its genetic modifiers. Assuming that the polygenic architecture of brain variation is driven by a mixture of molecular and cellular processes that have higher or lower contribution in a region-specific manner, we suspect that ICA may be able to tease apart, to some extent, these biological processes into less heterogeneous (and therefore more interpretable) genomic drives. A typical multi-modal MRI dataset contains thousands of regional traits or millions of voxel-wise traits, thereby yielding a large amount of genome-wide association data. By mining such data, we observed that loci in distant parts of the genome have concerted influences on brain traits, indicating that ICA captures variants and, indirectly, gene effects that potentially act within the same biological pathways and molecular mechanisms. Our observation that a number of highly polygenic components sensitively map to distinct molecular pathways supports this notion.

We observed different genetic underpinnings for various MRI modalities and contrasts. This can be of relevance to MRI biomarker discovery and imaging genetics, as it implies that each imaging modality captures a different aspect of neurobiology from a different angle. Similar to the previous research^9^, we observed that the genetic modifiers of cortical thickness and surface area are divergent and partly opposing. In contrast, different diffusion MRI metrics showed more overlap in their genetic underpinnings regardless of the data analysis methods used, including tractography and tract-based spatial statistics. This observation supports the notion that deeper insight can be gained by combining data across modalities, as modalities seem to complement each other in the space of genetic drives (Figure 3). In addition, taking advantage of multivariate patterns of effects can increase power, because probabilistic ICA enables modelling (and therefore removing) random noise^13^ that compounds the individual trait-GWASs.

Genomic sources can readily be applied to individual-level genotype information. By developing a universal genomic ICA atlas of our latent sources in a large population, the polygenic risk of each individual can be partitioned into independent (population-common) components using linear regression, similar to the ICA-based dual-regression method in functional neuroimaging^27^ and to the generation of polygenic risk scores in genetics^28^. Calculating loadings of different genomic sources by this strategy may inform about the genetic profile of an individual in relation to a combination of different brain-related traits and conditions. In this application, genomic ICA may overcome some of the current challenges in the translation of GWAS results in heterogeneous traits and populations. The ultimate aim of our approach will be to stratify individual patients with psychiatric and/or neurological conditions along the dimensions of the genomic sources, and understanding to what extent different biological processes are reflected in each genomic source potentially relevant for each individual. This can be a new strategy to segregate etiologic heterogeneity and moving towards precision medicine.

While we applied ICA to the genotype-phenotype associations observed in adult subjects, it is noteworthy that gene expression, DNA methylation and mRNA translation are regulated across cell types and often time-locked to developmental windows in brain networks. Considering that the polygenic risk of a diseases predisposes to pathology by altering the expression of critical gene sets at various stages, essential information may be extracted by the genomic source decomposition method. We provide preliminary evidence that different genomic sources align with the transcriptomic profile of different brain cell types, in both adult brain cells and in foetal brains. In the future, the same method can be applied to bimodal or multimodal data such as GWAS and single-cell transcriptomes (TWAS). Also, the time variable may be included in the model as an extra dimension along which gene expression changes may be decomposed, for example using temporal ICA^29^ or tensorial ICA^30^. Such methods can disentangle disease factors into mechanistically coherent processes that are specific to certain cell types, developmental windows and polygenic variants. This strategy may also improve our understanding of the MRI biomarkers, modalities and their cell-type specificity.

A limitation of our work is the arbitrary choice of 200 components for source separation. The choice of ICA dimensionality, or *model order estimation*, is a topic of ongoing research for which no consensus exists^31^. Various methods are used for this purpose, such as the maximum likelihood criterion, Akaike’s information criteria (AIC) or minimum descriptive length (MDL^6^). However, these methods make assumptions about the raw data and the goodness of fit that may not align with the nature of the genome-wide data in practice. One workaround is a brute-force-like application of ICA at many dimensions and choosing the decomposition that achieves the highest performance with regard to some biological measures(s) of interest *post hoc*. These measures may include correlation of the genomic components with individuals’ behavioural or neuropsychiatric phenotypes.

In summary, we propose genomic ICA as a new method to identify latent genetic factors that influence brain structure and function as measured using MRI. The derived latent dimensions are highly sensitive to known molecular pathways and cell-specific gene expression data. In this way, genomic ICA reduces data produced by many univariate GWASs in a way that aids mechanistic interpretation. Thus, genomic ICA may help to disentangle the many different biological routes by which the genome drives inter-individual variation in the brain. Future research is aimed at further streamlining and validating genomic ICA, generalising its application to other types of *omics* data, and evaluating its clinical utility in biologically driven stratification of heterogeneous patient populations.

## Acknowledgements

This work was supported by a Radboudumc Hypatia grant (to Emma Sprooten), BBRF, Christine Mohrmann. MPI. UK Biobank and Oxford BIG. SurfSara: This work is part of the research programme *Computing Time National Computing Facilities Processing Round pilots 2018* with project number *17666*, which is (partly) financed by the Dutch Research Council (NWO) and was carried out on the Dutch national e-infrastructure with the support of SURF Cooperative.

## Supplementary Materials

[**Supplementary figure 1:** Manhattan plots of all 200 components].

**Supplementary figure 2.**
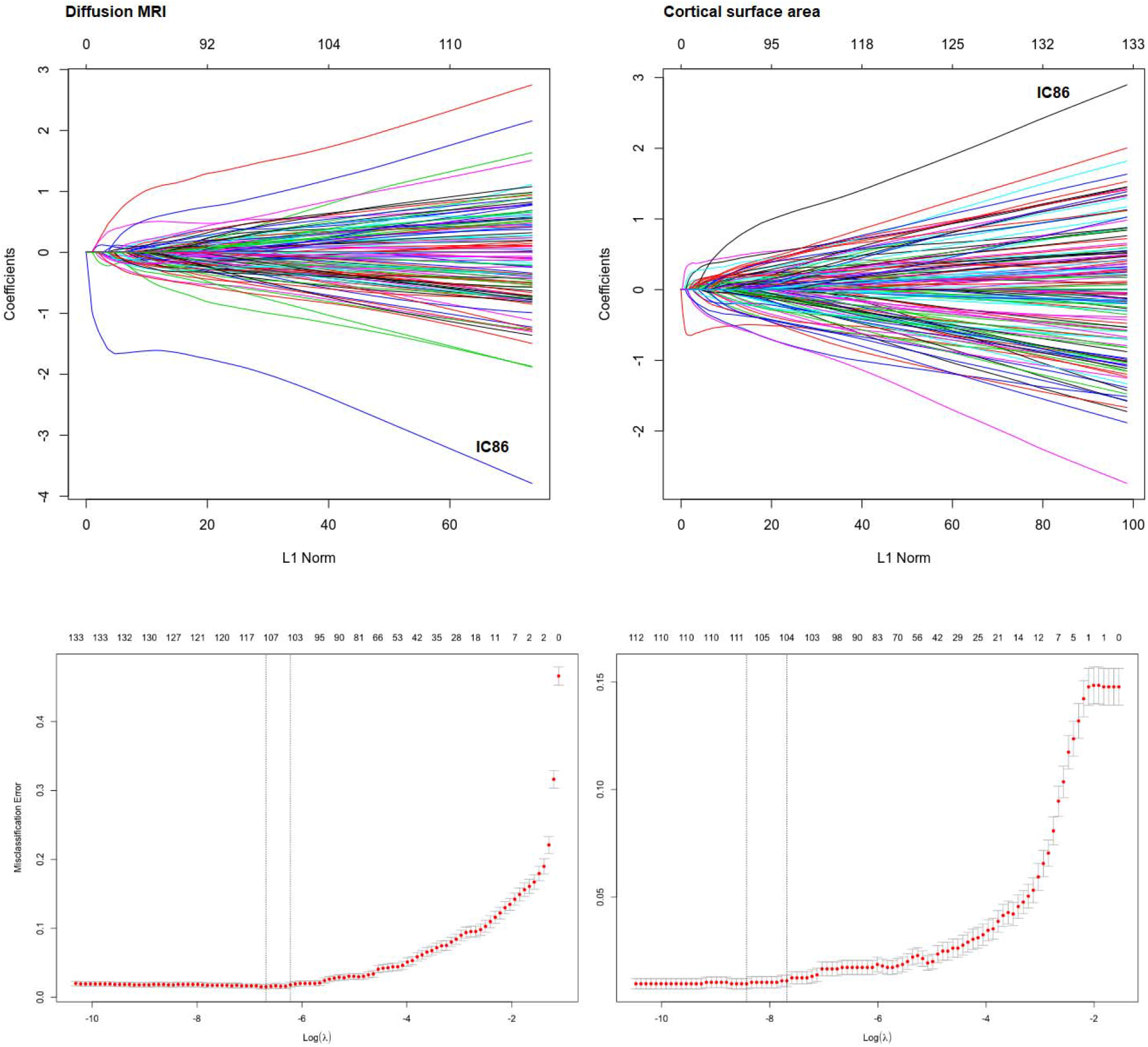
Genomic components predicting diffusion and cortical surface area features. Logistic regression using L1 regularization (LASSO) highlights a number of components with higher predictive power, including IC86 as the top predictor for both modalities.

