## Supplementary figure 1 for "Independent Genomic Sources of Brain Structure and Function"

Supplementary figure 1. Genomic component Manhattan plots

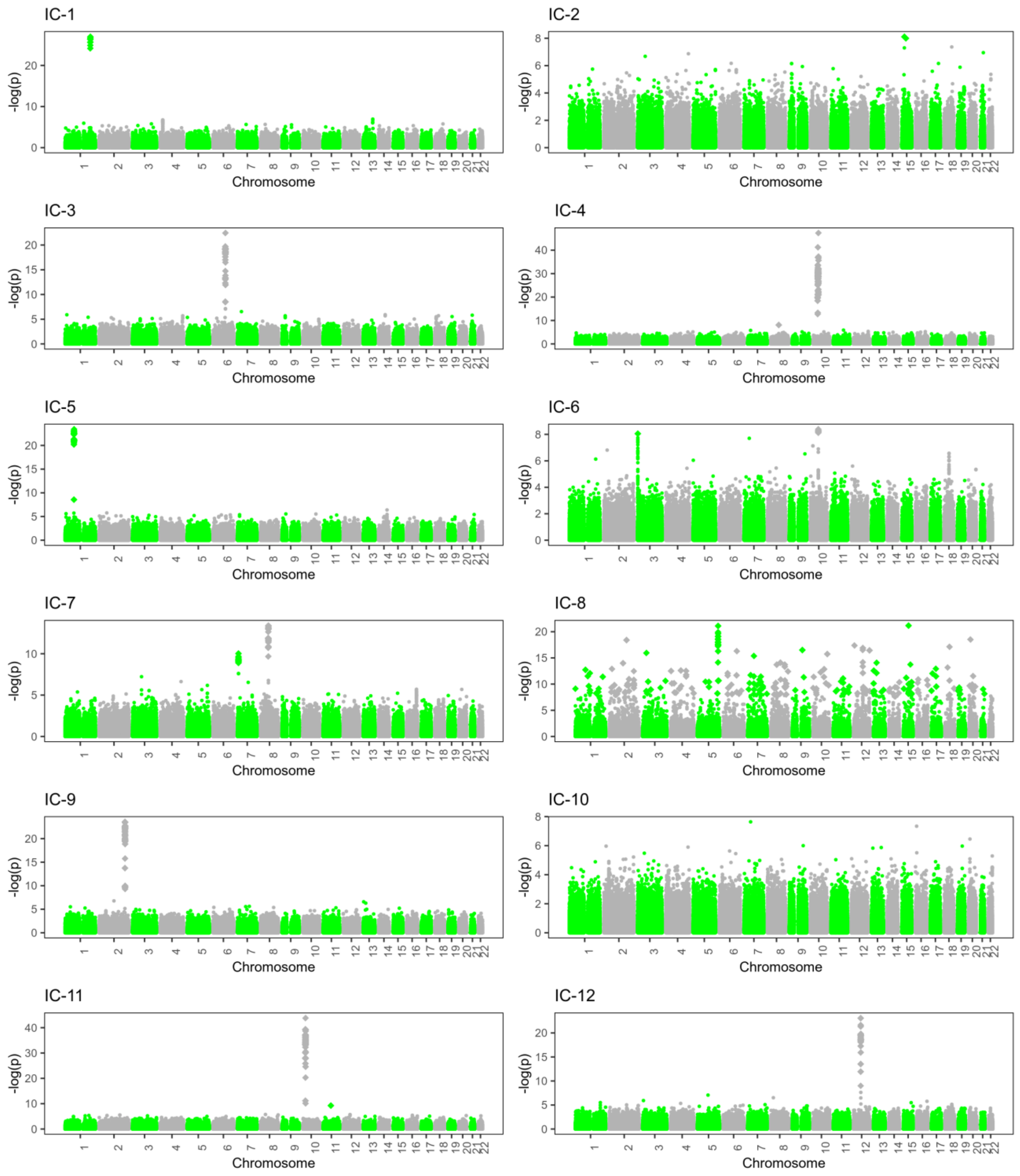

Supplementary figure 1. Genomic component Manhattan plots

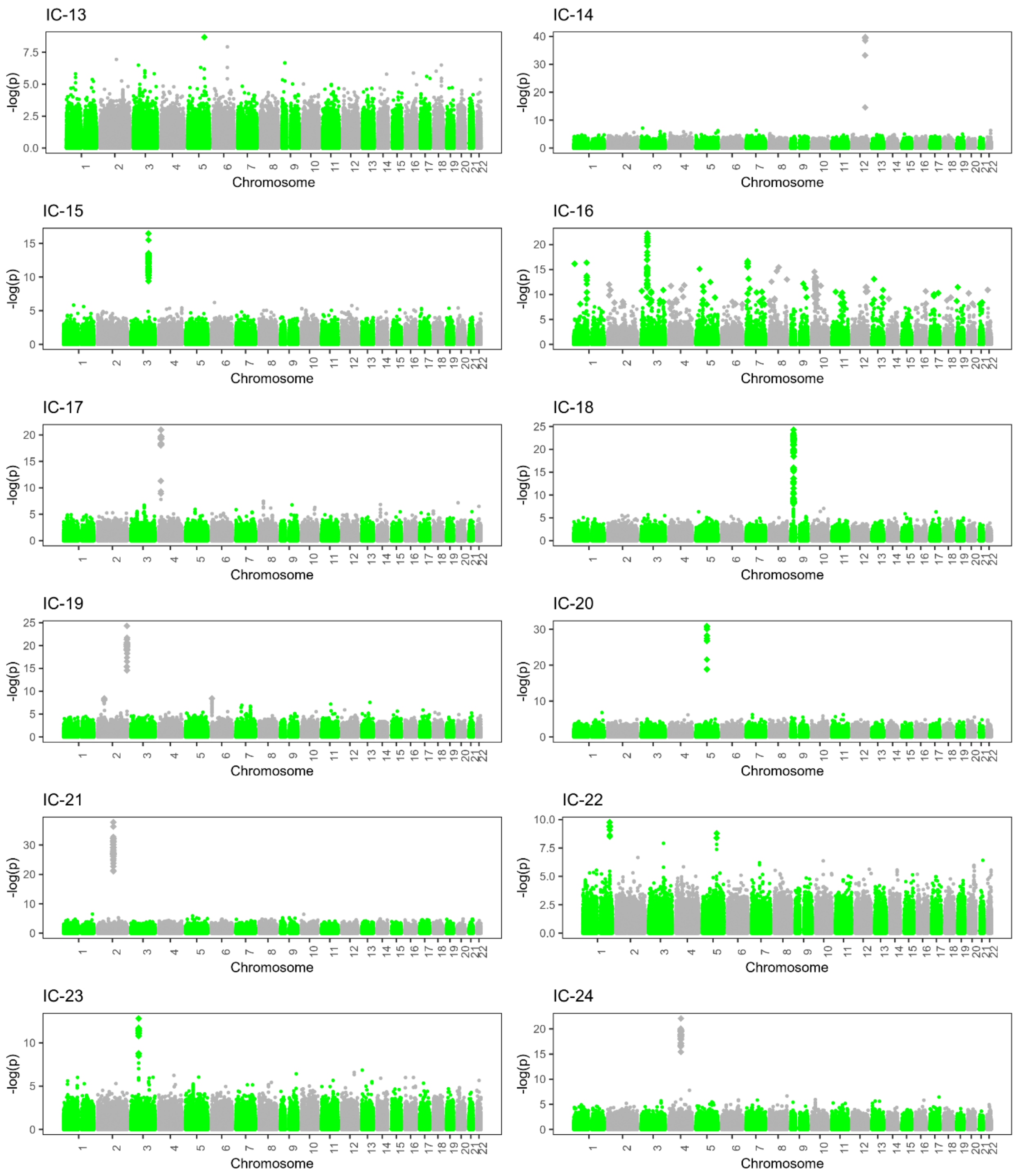

Supplementary figure 1. Genomic component Manhattan plots

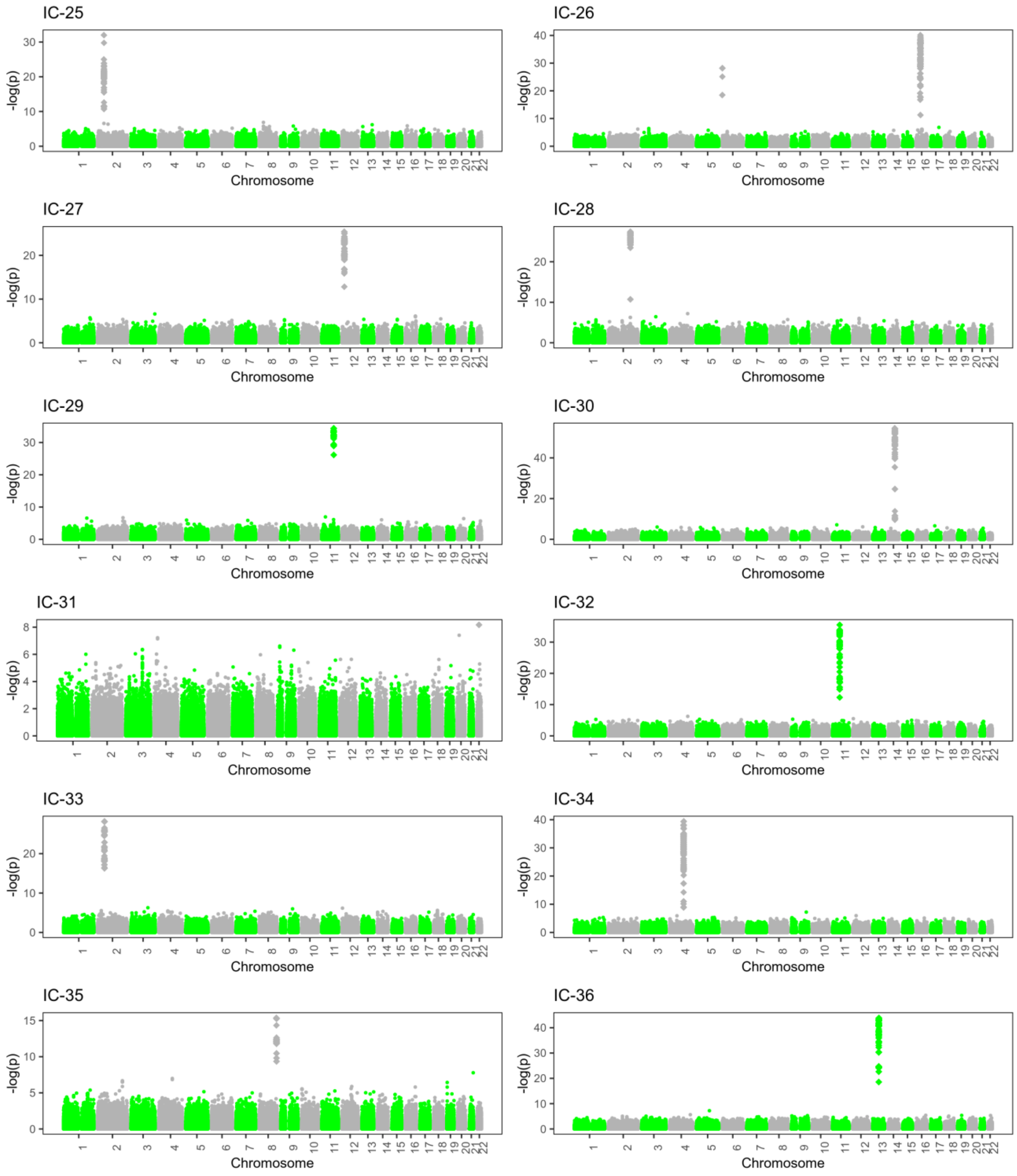

Supplementary figure 1. Genomic component Manhattan plots

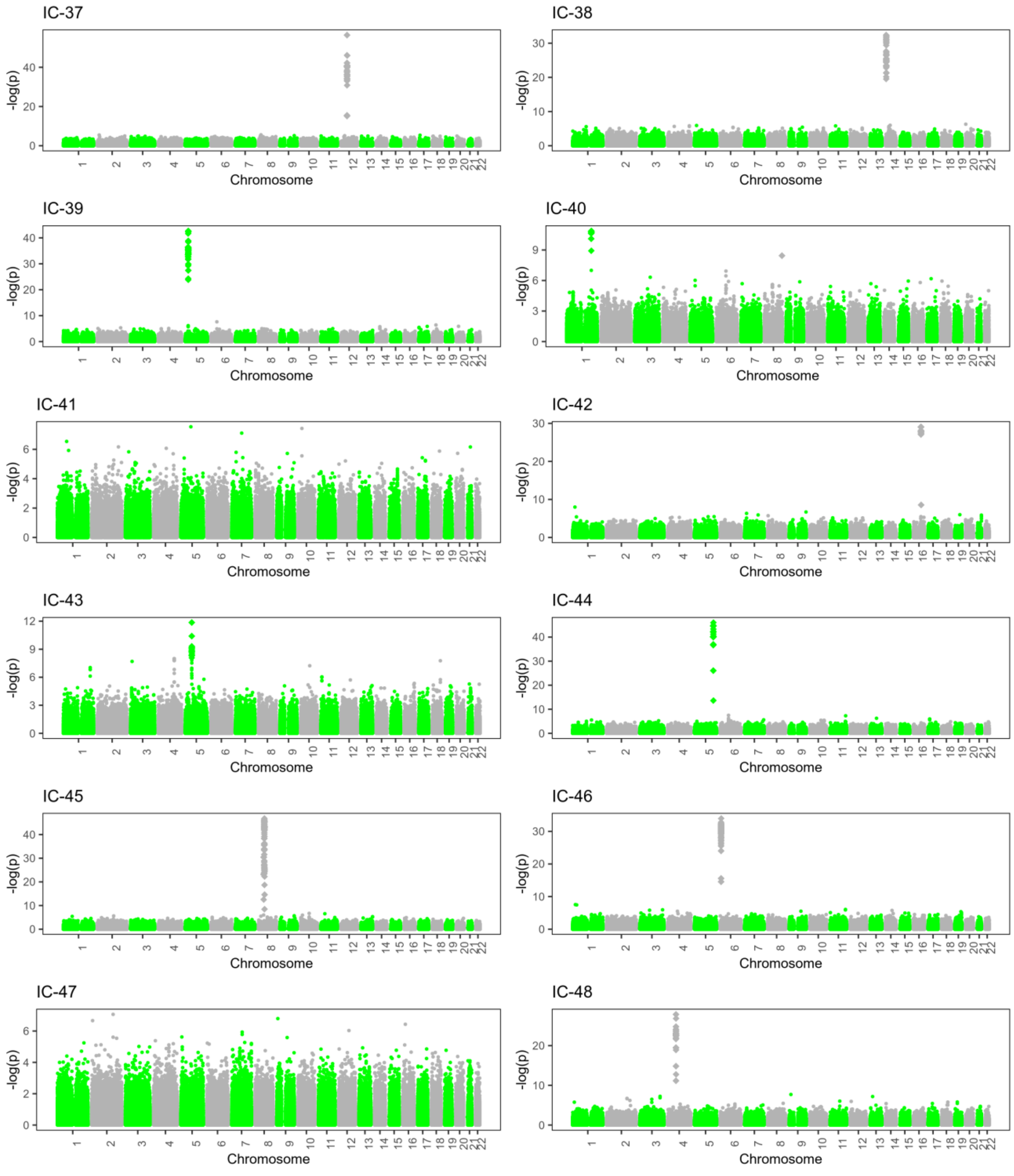

Supplementary figure 1. Genomic component Manhattan plots

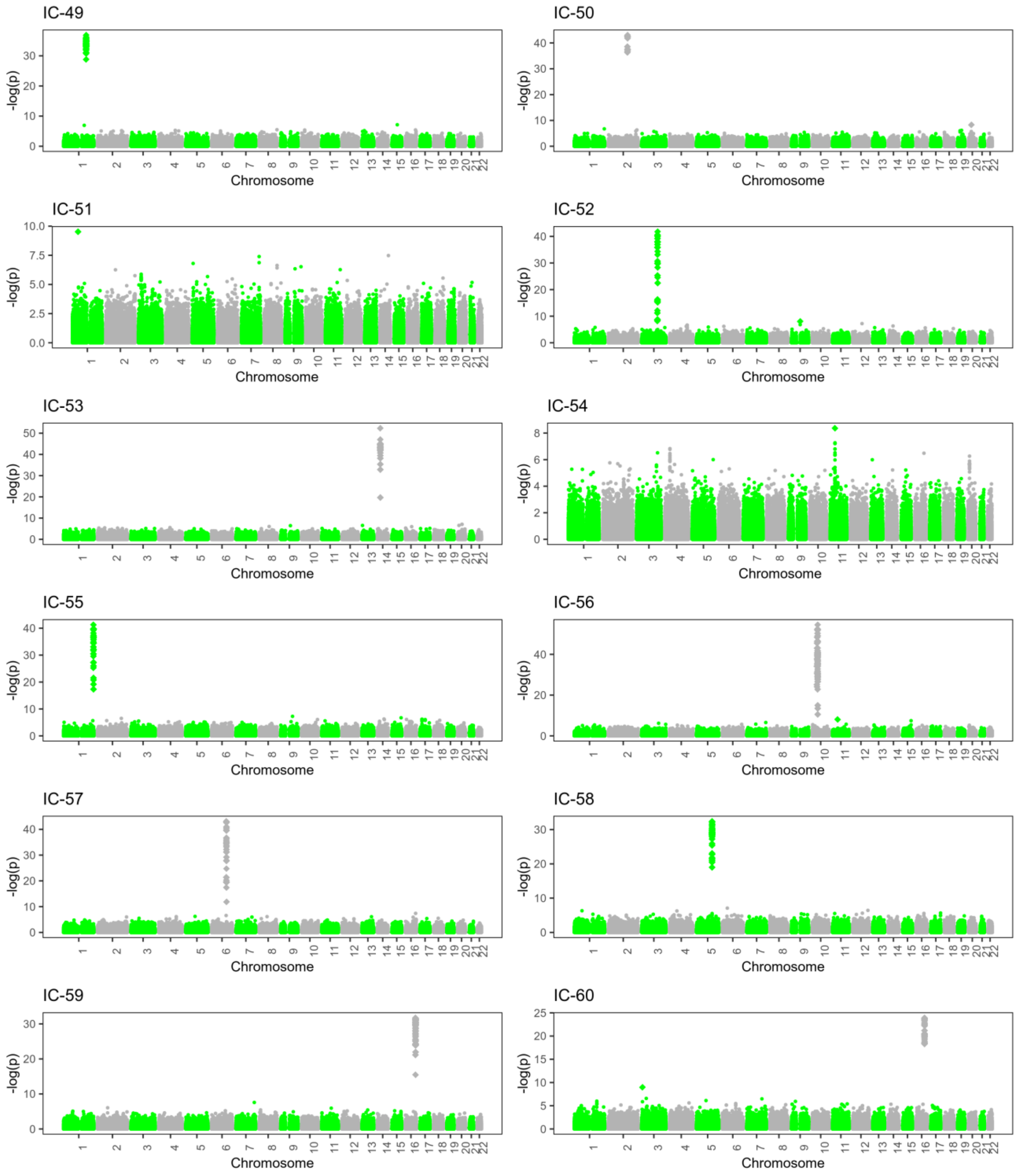

Supplementary figure 1. Genomic component Manhattan plots

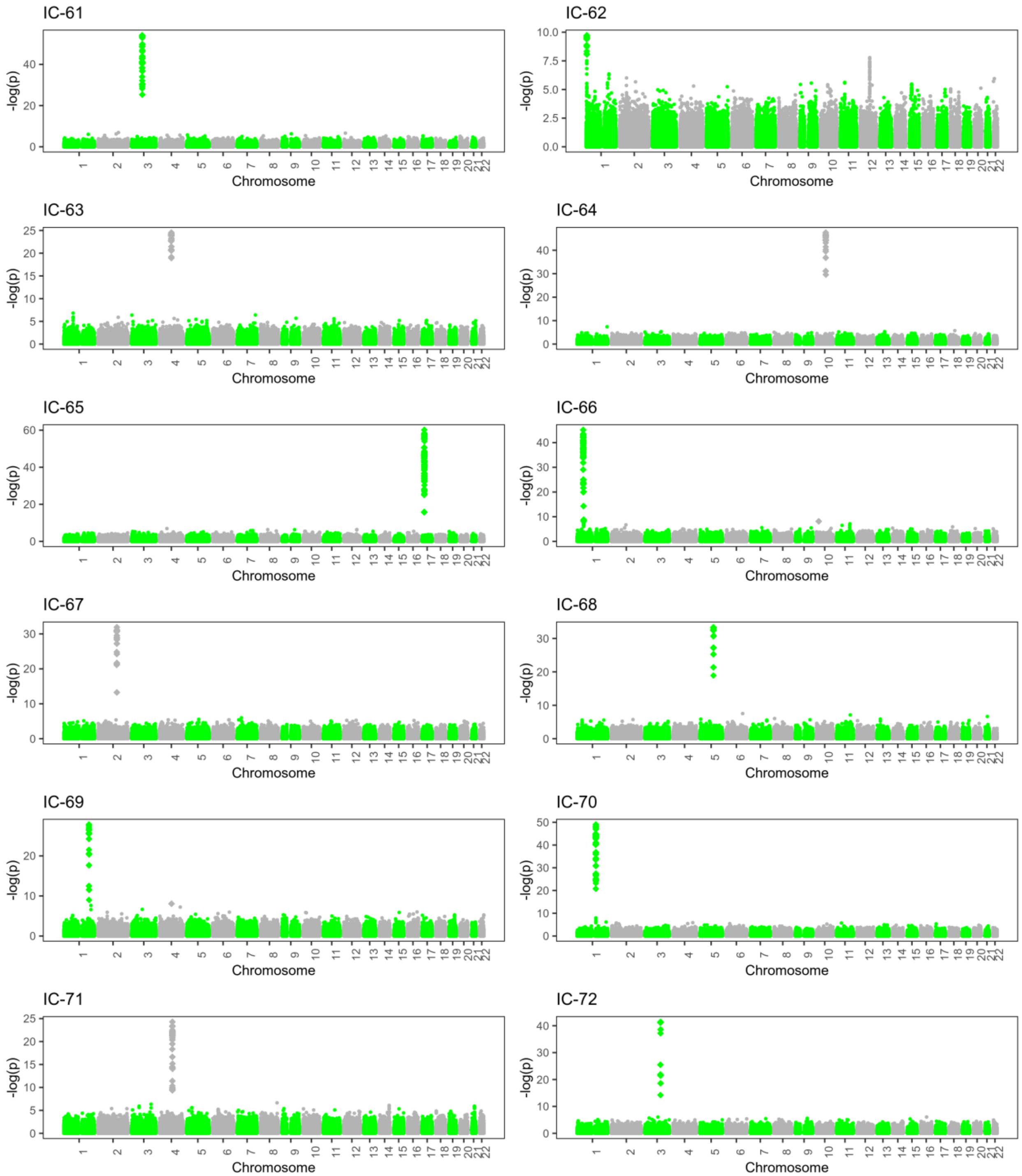

Supplementary figure 1. Genomic component Manhattan plots

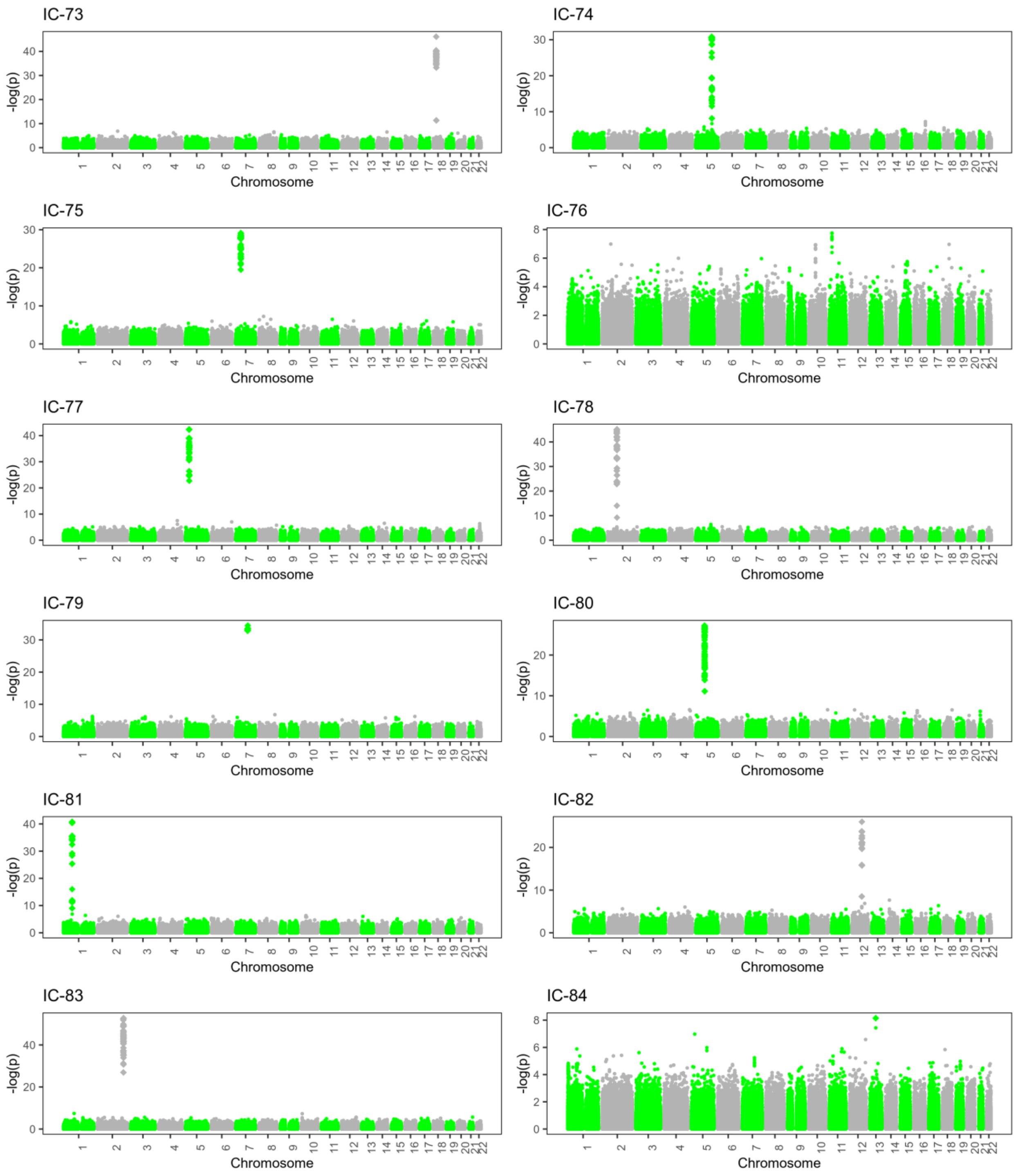

Supplementary figure 1. Genomic component Manhattan plots

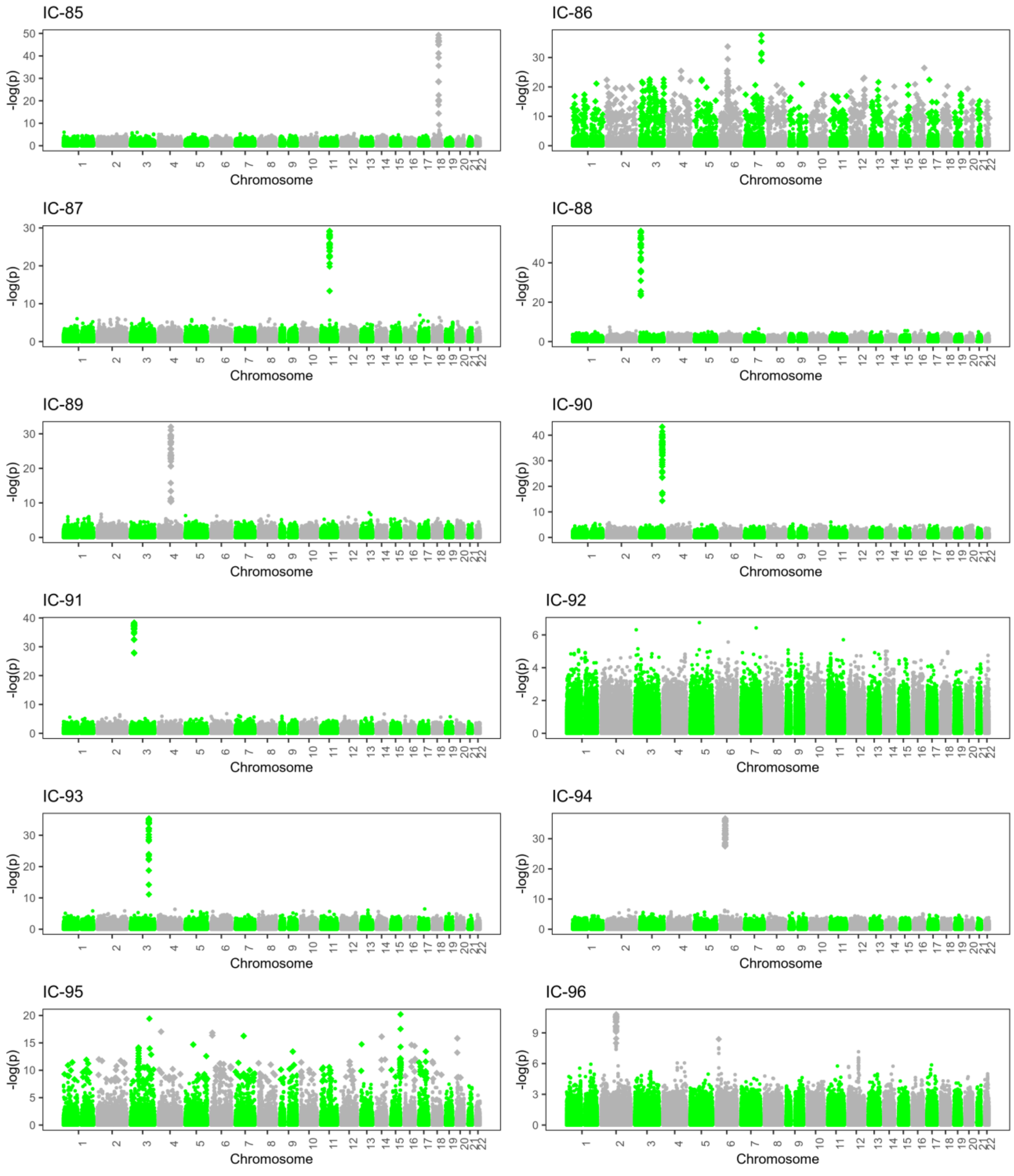

Supplementary figure 1. Genomic component Manhattan plots

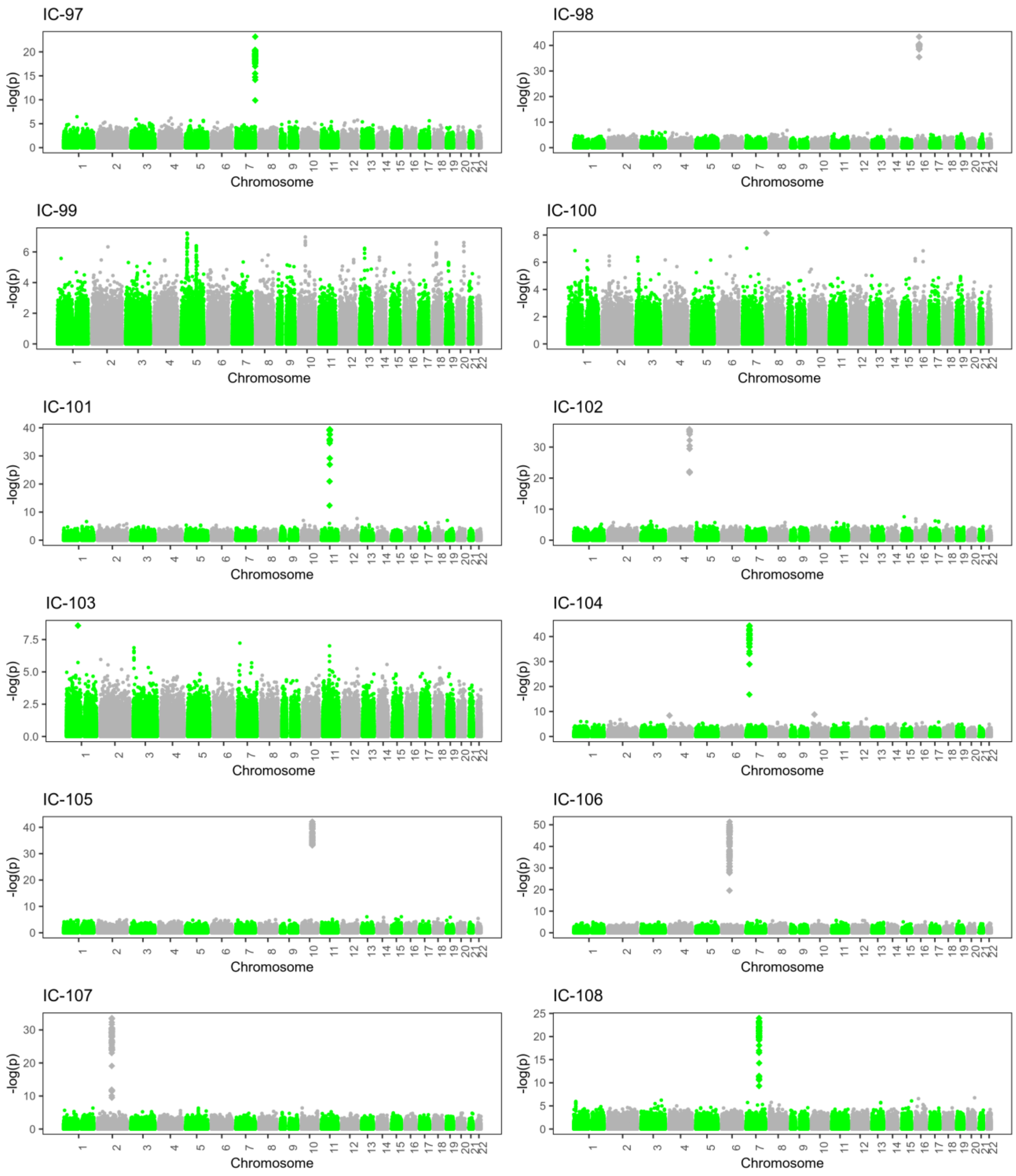

Supplementary figure 1. Genomic component Manhattan plots

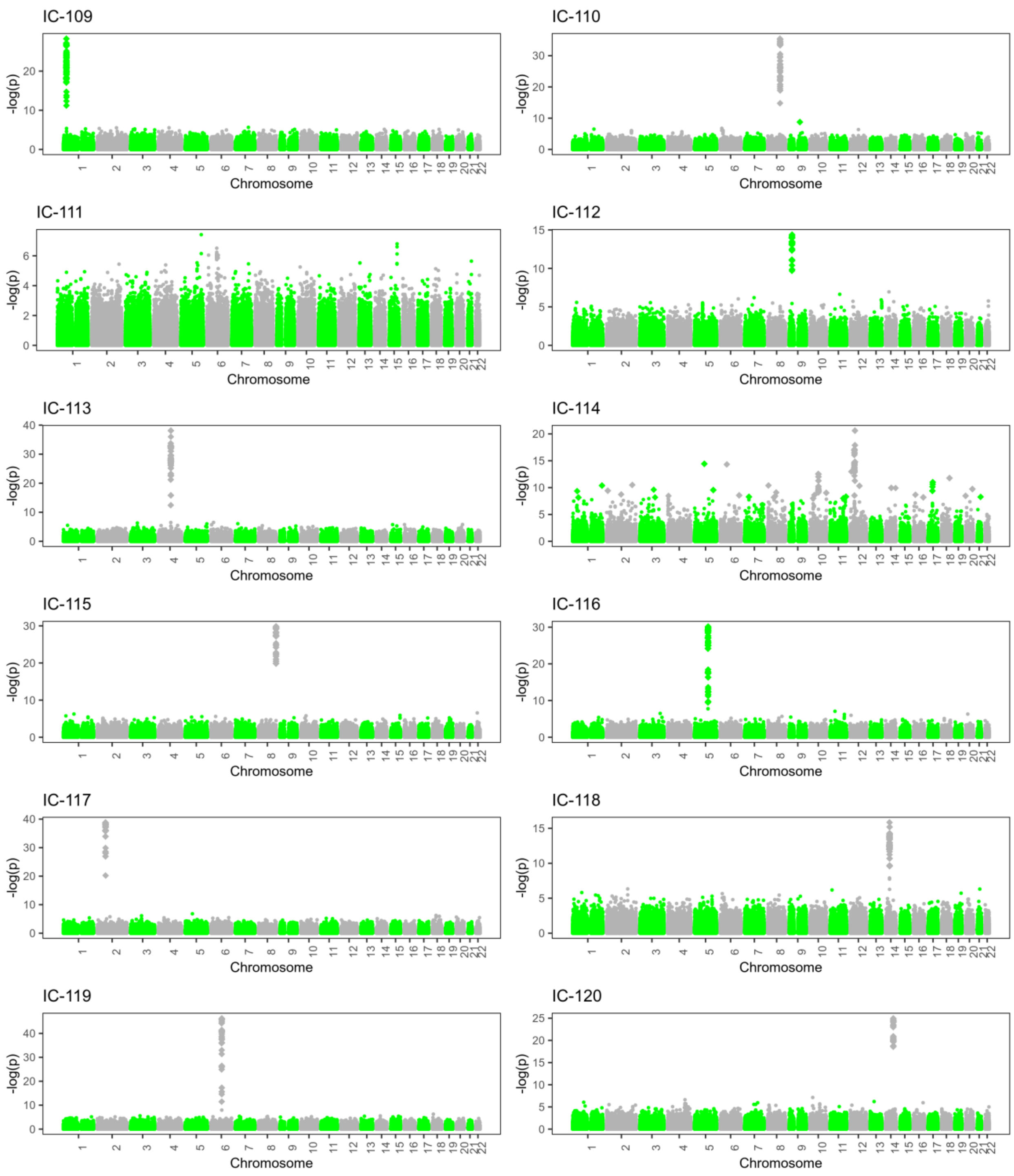

Supplementary figure 1. Genomic component Manhattan plots

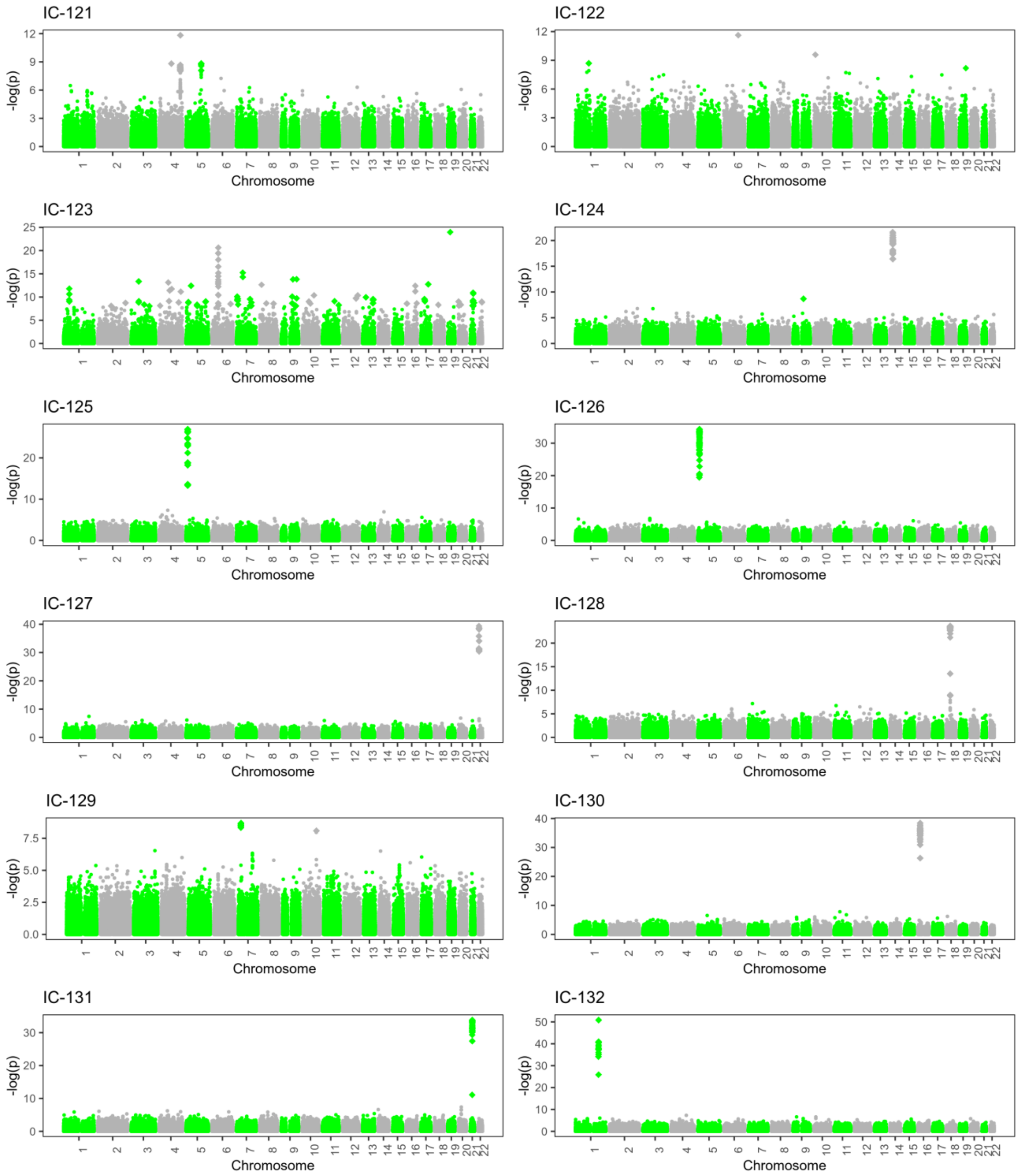

Supplementary figure 1. Genomic component Manhattan plots

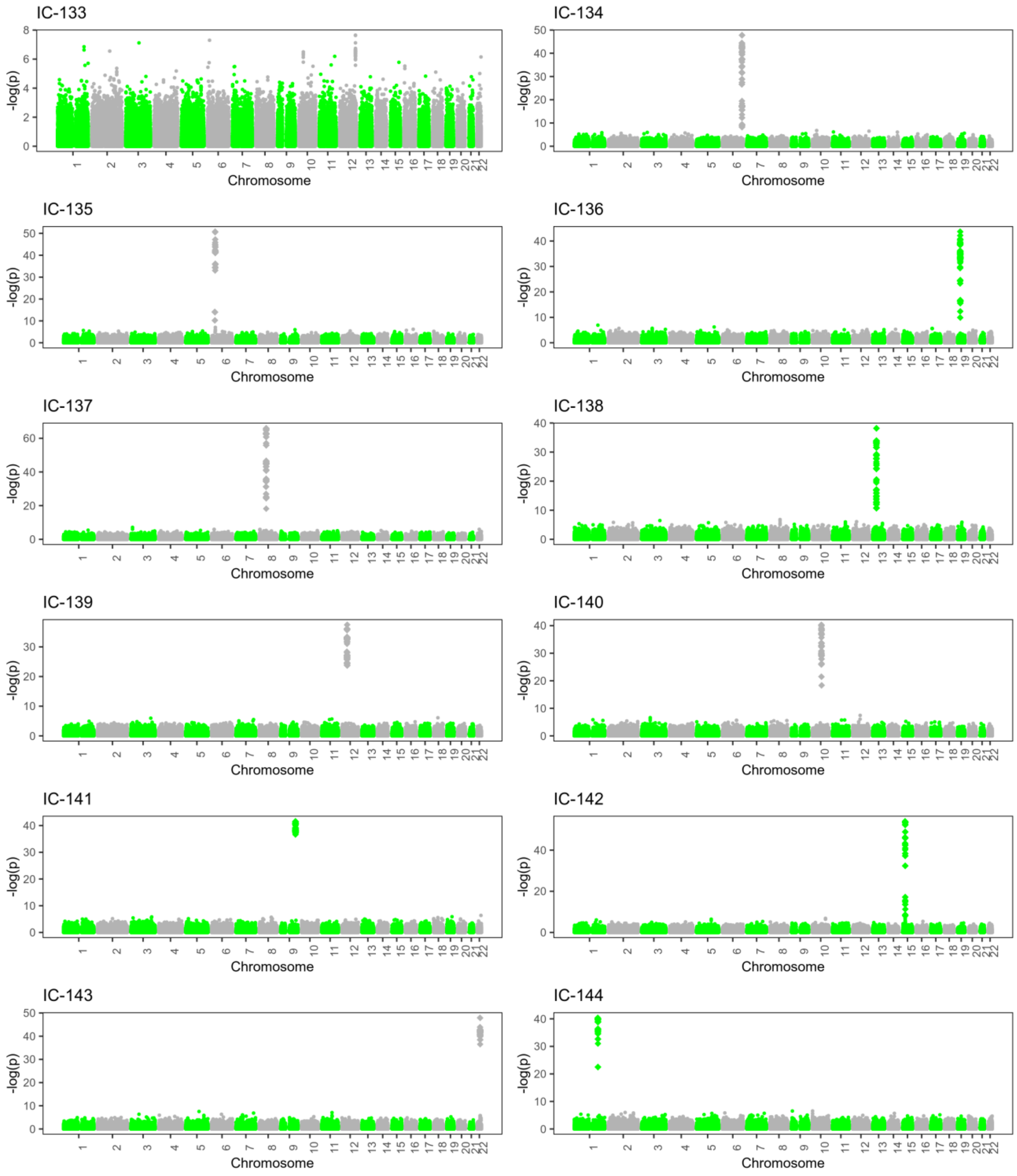

Supplementary figure 1. Genomic component Manhattan plots

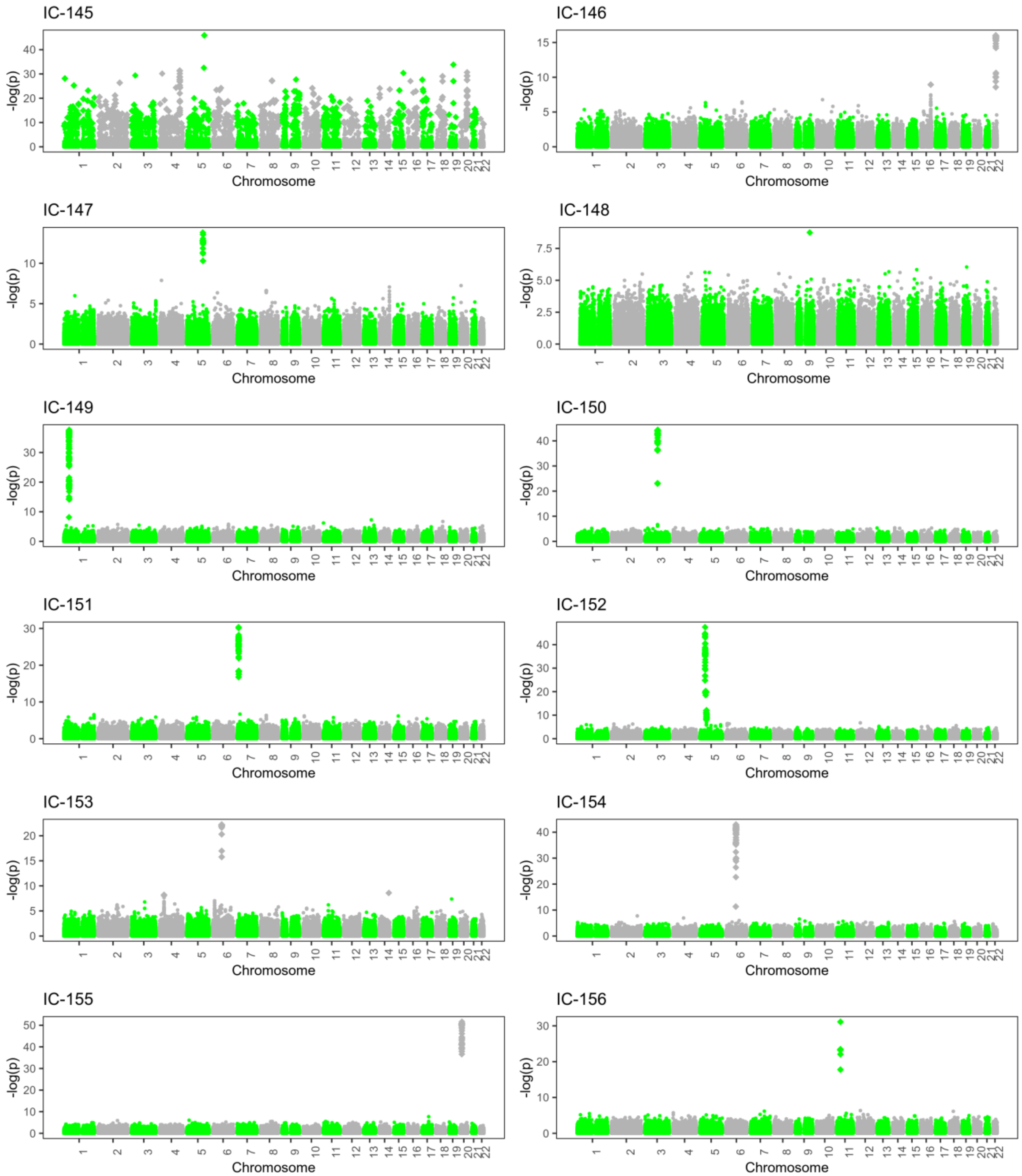

Supplementary figure 1. Genomic component Manhattan plots

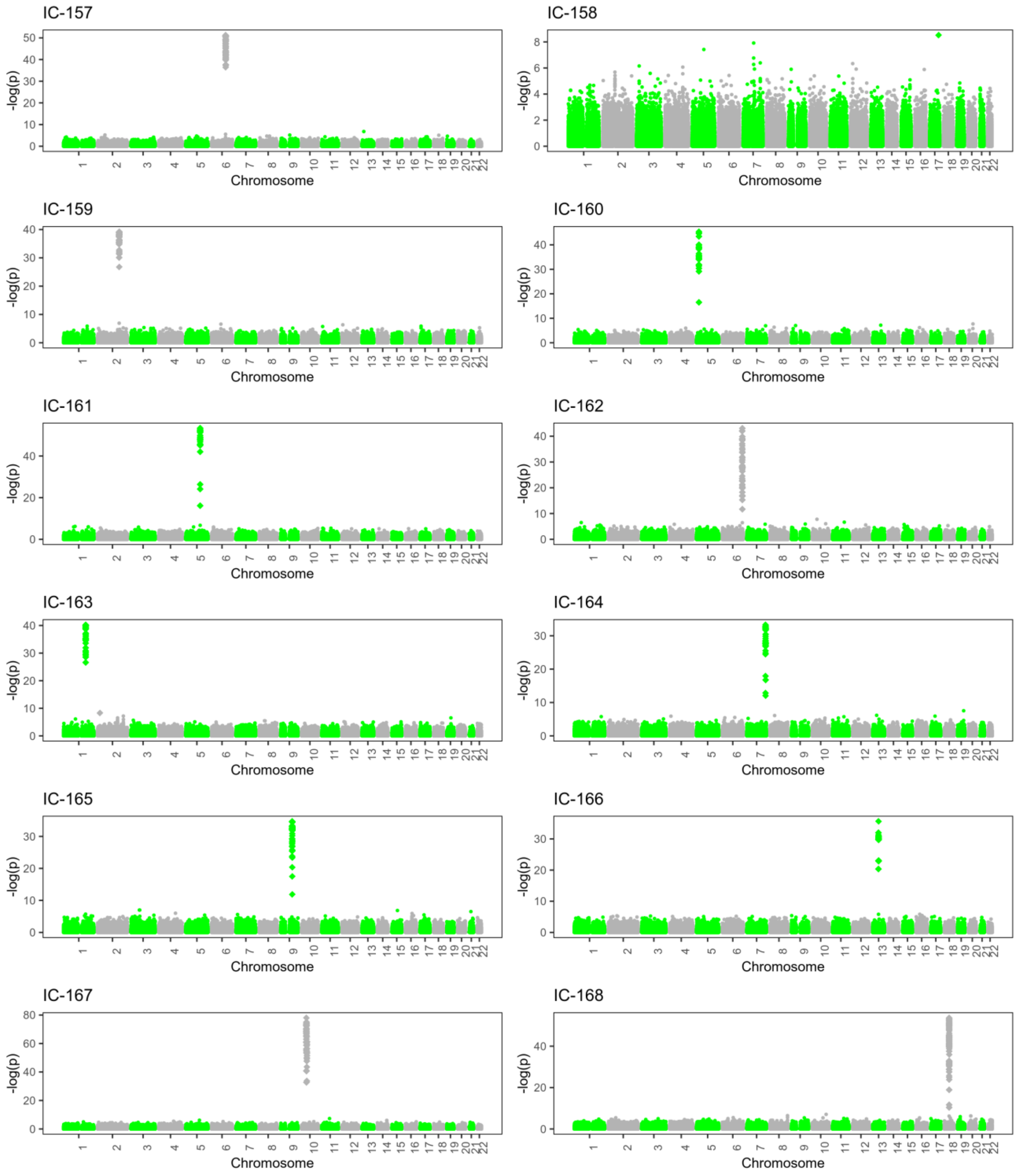

Supplementary figure 1. Genomic component Manhattan plots

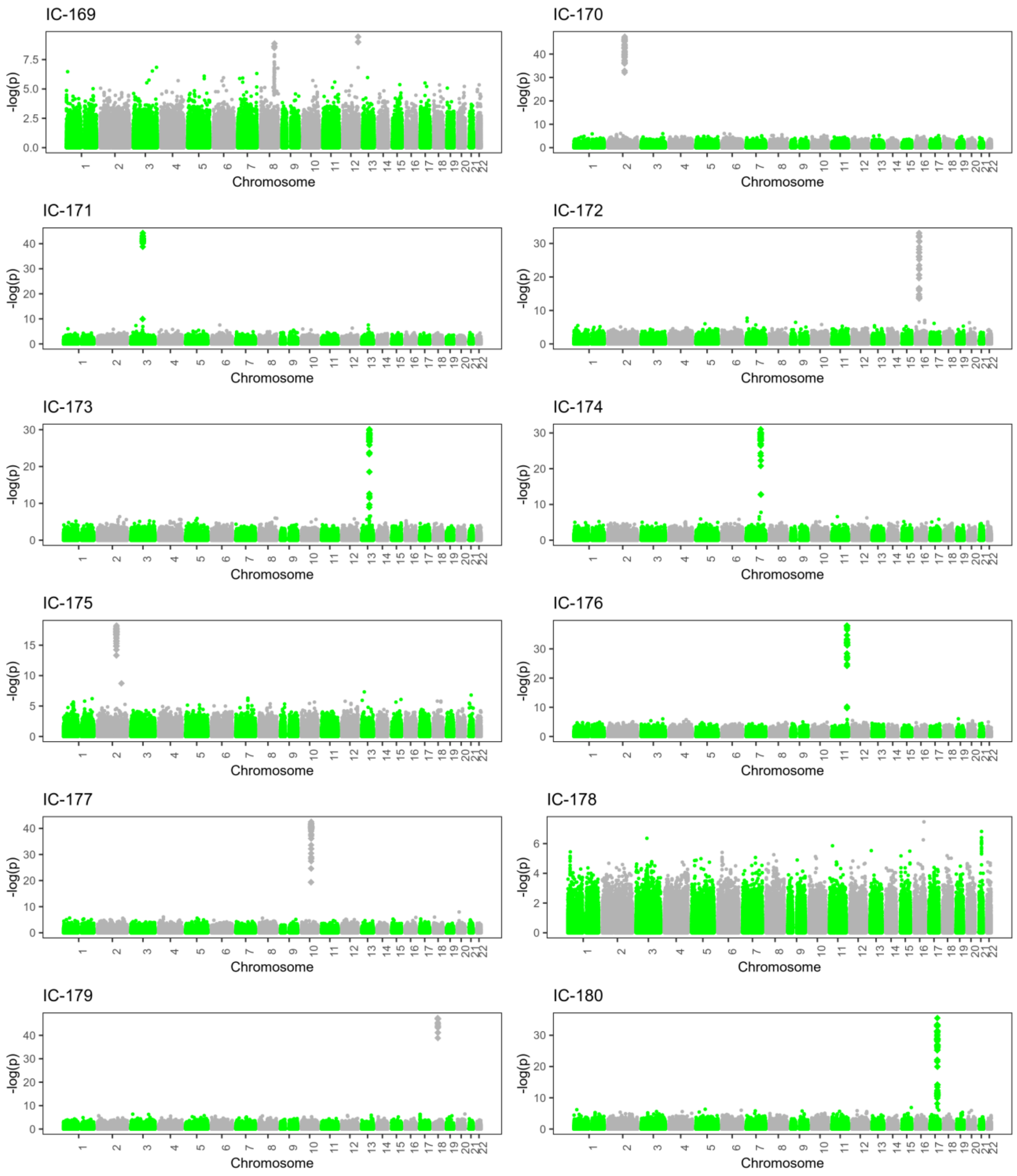

Supplementary figure 1. Genomic component Manhattan plots

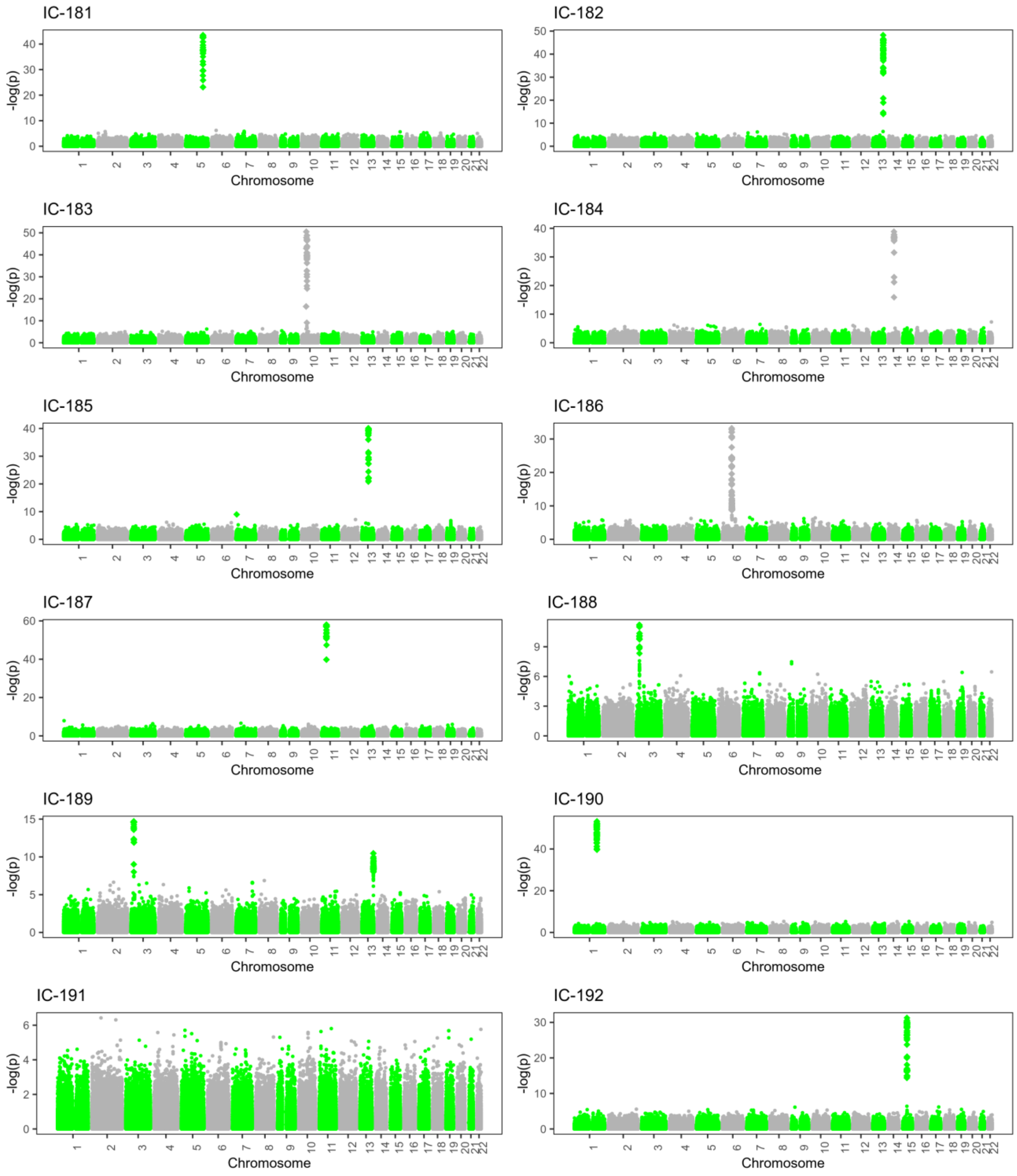

Supplementary figure 1. Genomic component Manhattan plots

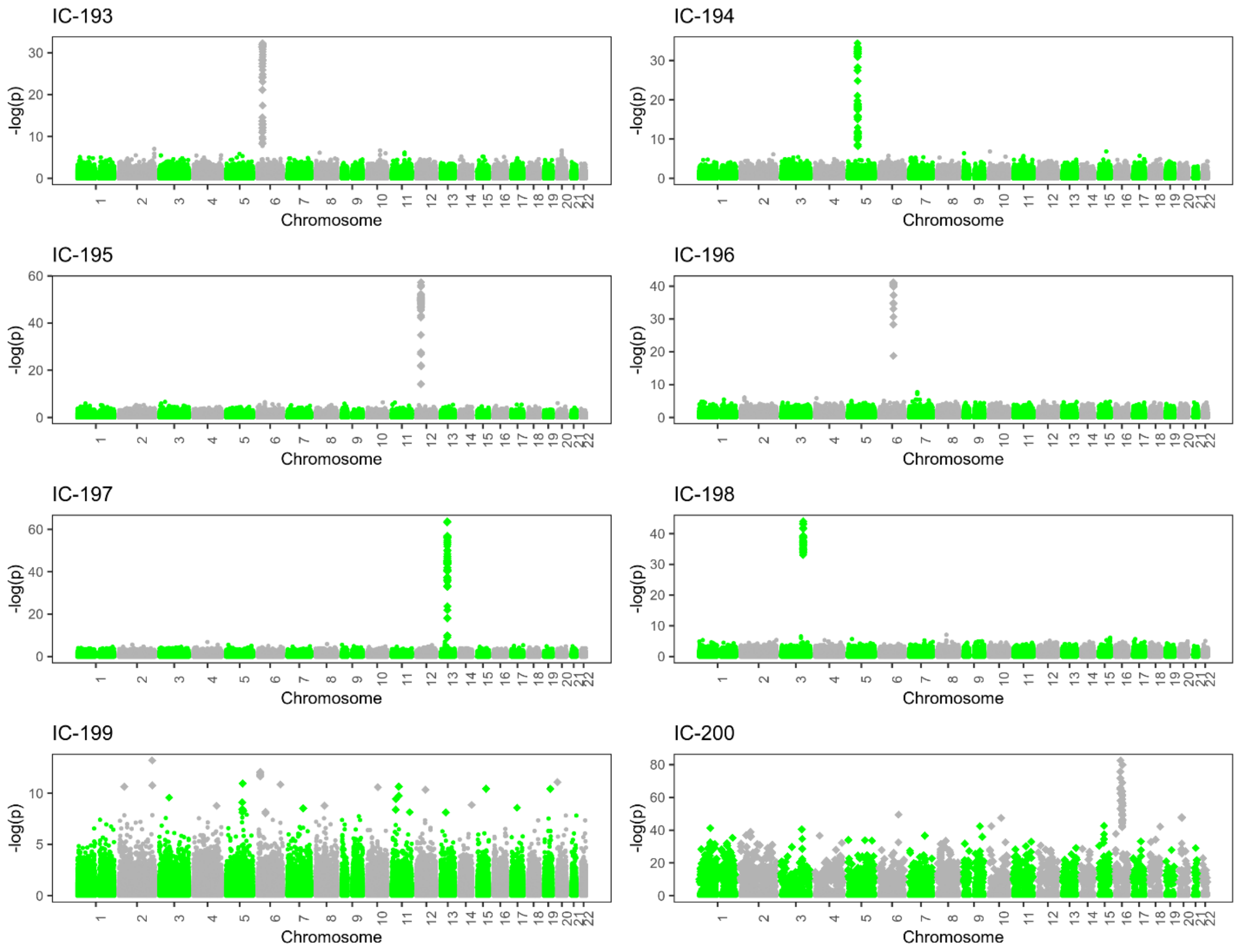
